# Coarse-Grained Simulations of Mycobacterial Outer Membranes Reveal Fluidity-Dependent PDIM Redistribution Across Different Lipid Environments

**DOI:** 10.64898/2026.02.18.706594

**Authors:** Bibek Acharya, Sudarshan Lammichane, Turner P. Brown, Matthieu Chavent, Wonpil Im

## Abstract

The mycobacterial outer membrane (MOM) constitutes an asymmetric permeability barrier that influences lipid organization and transport in *Mycobacterium tuberculosis*. In this study, we have developed MARTINI 3 coarse-grained (CG) lipid models of the MOM, incorporating α-mycolic acids, 5 different trehalose-based lipids, and PDIM (phthiocerol dimycocerosate). The CG models were parameterized and validated using all-atom simulations of symmetric inner- and outer-leaflet membranes, as well as fully asymmetric MOM models. Bonded parameters were optimized through an iterative refinement procedure targeting atomistic bonded distributions. The CG simulations show good agreement with the all-atom simulation data and available experimental measurements in terms of the membrane thickness, solvent accessible surface area, lipid density profiles, and the outer-leaflet-induced lipid disorder in α-mycolic acids at the inner leaflet. The model reproduces the temperature-dependent phase behavior of all-atom α-mycolic acid membranes. Using this model, we demonstrate that PDIM localization, diffusion, and aggregation are strongly modulated by membrane fluidity and lipid composition, with enhanced translocation and clustering in liquid disordered environments. Our CG MOM lipid models provide a validated platform for large-scale simulations of mycobacterial membranes and enable mechanistic studies of lipid organization, membrane dynamics, and protein-membrane and membrane-drug interactions.

## Introduction

*Mycobacterium tuberculosis* (*Mtb*), the etiological agent of tuberculosis (TB), is a slow-growing, acid-fast bacterium that primarily infects the lung and affects other parts of the body^1,2^. Despite the existence of antibiotics and vaccines, TB continues to affect 10 million individuals annually with more than a million annual mortalities, making itself one of the most consequential, infectious diseases globally and historically^3^. The global burden of TB is further compounded by the emergence of multidrug-resistant (MDR) and extensively drug-resistant (XDR) strains, which not only complicate diagnosis and treatment, but also undermine current therapeutic strategies^4,5^. While the cell envelope of *Mtb* is a common target of anti-TB drugs, emerging evidence suggests that *Mtb* can adapt by altering the composition of its cell envelope in response to its environment or drug-induced stress^6–8^. Although several studies have investigated the evolution of the *Mtb* cell envelope over time, a significant knowledge gap remains regarding how these structural changes are linked to the development of drug resistance.

The *Mtb* cell envelope is a thick, complex, and multilayered structure that serves as a protective barrier, enabling the bacterium to survive in diverse and often hostile environments^9,10^. At its innermost level lies the plasma membrane (or mycobacterial inner membrane), typical of bacterial membranes, consisting of phospholipids, phosphatidyl-*myo*-inositol mannosides (PIMs), lipomannan, and lipoarabinomannan. Surrounding this is a robust cell wall core consisting of peptidoglycan covalently linked to arabinogalactan, which is further esterified to be linked to long-chain (C60-C90) mycolic acids (MAs). These covalently bound MAs constitute the inner leaflet of the mycobacterial outer membrane (MOM) or mycomembrane, while a diverse array of noncovalently associated lipids and glycolipids, including phthiocerol dimycocerosate (PDIM), trehalose-based lipids such as trehalose dimycolate (TDM), trehalose monomycolate (TMM), diacyltrehalose (DAT), pentaacyltrehalose (PAT), and sulpholipid (SGLs), forms the outer leaflet of the MOM. Collectively, these MOM components give rise to the highly asymmetric and hydrophobic nature, which accounts for approximately 60% of the cell envelope’s dry weight. This unique and impermeable lipid bilayer plays a pivotal role in *Mtb* drug resistance, immune evasion, and host-pathogen interactions. The outer leaflet, enriched with highly hydrophobic, *Mtb*-specific lipids described above, substantially contributes to the organism’s resilience and limited membrane permeability^7,9–13^.

The unusual lipid diversity and organization of the MOM pose major challenges for experimental characterization. As a result, computational approaches such as molecular dynamics (MD) simulations have become indispensable for probing lipid organization, membrane dynamics, and drug interactions at high resolution. While all-atom (AA) MD simulations^14–21^ provide valuable insights into the behavior of specific mycobacterial lipids and small membrane patches, their applicability remains limited by the system size and computational cost. These constraints hinder our ability to capture the full structural complexity and longer timescale dynamics of the MOM. Therefore, there is a growing need for validated multiscale simulation frameworks that combine AA accuracy with coarse-grained (CG) efficiency, enabling the study of large, compositionally realistic MOM systems under diverse physiological and pharmacological conditions^22–24^.

Despite the advantages of CG modeling, comprehensive and validated CG lipid models of the *Mtb* MOM are still lacking. Existing studies focus on individual lipids or simple bilayers, without incorporating the full complexity of the *Mtb* lipidome^25,26^. To fill this gap, in this study, we present a MARTINI 3 CG model of the *Mtb* MOM that includes the major lipid constituents (α-MA, PDIM, SGL, PAT, DAT, TMM, and TDM) identified in experimental studies. We have validated the model by comparing bonded distributions, solvent accessible surface area (SASA) of individual lipids, global membrane properties such as bilayer thickness, and lipid density distributions against AA simulations and available experimental data. Our goal is to provide a robust and scalable platform for studying the structural and dynamic heterogeneity of the *Mtb* MOM under various physiological conditions and to facilitate future studies on drug permeability, protein-lipid interactions, and membrane-targeting therapies for TB.

## Methods

### System Building and Simulation

For CG model parameterization and validation, we used the following AA simulations from the recent work by Brown et al^15^: (1) symmetric MOM inner-leaflet systems used to characterize the α-MA bilayers, (2) symmetric MOM outer-leaflet systems used to characterize the mixed bilayers of PDIM and trehalose-based glycolipids (SGL, PAT, DAT, TMM, and TDM), and (3) asymmetric MOM systems created to combine the inner and outer leaflets into a single more realistic asymmetric MOM. The corresponding CG initial structures were generated by mapping the AA coordinates using our custom Python scripts. For the asymmetric MOM system, we also simulated 4X and 16X larger systems (in the membrane area, i.e., the xy plane) to check their stability and consistency with the 1X system. In addition to these systems, various single-component bilayers (POPC (palmitoyl oleoyl phosphocholine), PEPC (palmitoyl erucoyl phosphatidylcholine), PSPC (palmitoyl stearoyl phosphocholine) were used to investigate PDIM behavior in different lipid environments at different temperatures. These systems were built using in-house CHARMM-GUI Membrane Builder with 0.15M KCl^27–30^. The lipid compositions and the simulation information of all systems are summarized in **Tables S1-S2**.

All CG MD simulations were performed using GROMACS 2023.3^31,32^ following the parametrization rules recently updated for MARTINI 3 lipidome (**Figure 1**)^33^. Each system was equilibrated using a four-step equilibration protocol, in which the harmonic restraints on the lipid headgroup beads were gradually reduced at each step. This ensured proper packing of the bilayer before the production run. Production simulations were carried out in the NPT (constant particle number, pressure, and temperature) ensemble using a 20-fs timestep. Temperature was maintained using the velocity-rescale (V-rescale) thermostat^34^ with a coupling time of 1 ps, applied separately to the membrane and solvent. Pressure was controlled using the stochastic cell-rescale (C-rescale) barostat^35^ with a semi-isotropic coupling time of 4 ps and a compressibility of 3×10^−4^ bar^−1^ in both the lateral and normal directions. The electrostatic interactions were treated using the reaction-field method, and van der Waals interactions were treated using a potential-shifted Lennard–Jones scheme with a cutoff of 11 Å. Bond length constraints were maintained using the LINCS algorithm^36,37^. All production simulations were run for 10 μs with three replicas for each system unless explicitly mentioned (see **Table S2**). VMD^38^ and MartiniGlass^39^ were used for visualization.

**Figure 1.**
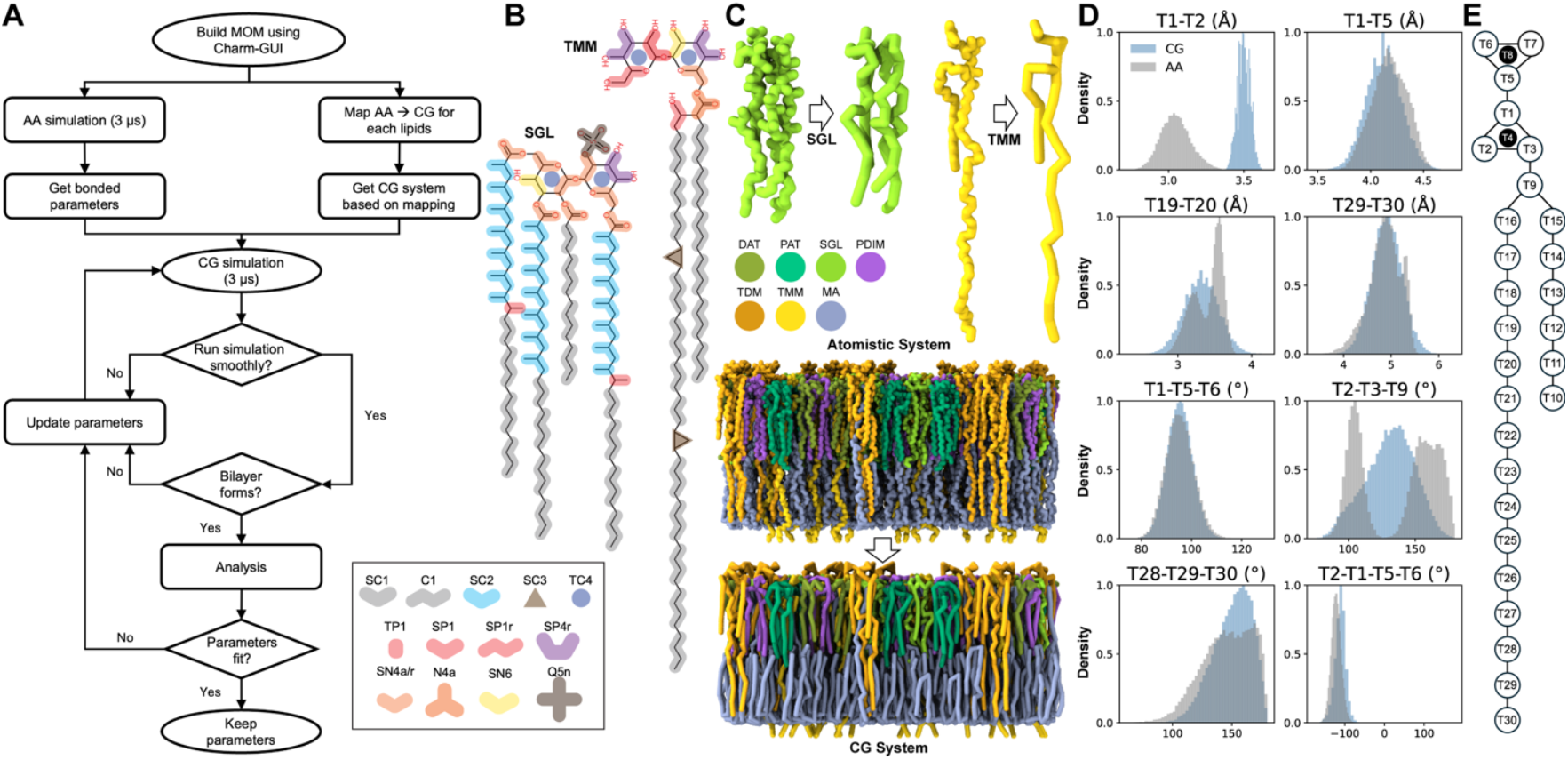
Development and parameterization of CG models for MOM lipids. **A**. Workflow for parameterizing *Mtb* outer membrane lipids from the AA to CG resolution. **B**. AA-to-CG bead mapping of SGL (left) and TMM (right) with the corresponding bead labels shown below them; see **Figures S1** for all lipids. **C**. AA and CG structure representations of SGL (top left) and TMM (top right), and the AA–CG mapped structures of an asymmetric MOM model (bottom) with lipid specific color codes. **D**. Bonded parameter optimization for the CG TMM model, including comparisons of bond lengths (two-bead definitions, e.g., T1–T2), bond angles (three-bead definitions, e.g., T1–T5–T6), and dihedral angles (four-bead definitions, e.g., T2–T1–5–T6) between AA and CG simulations. **E**. Bead naming scheme used for TMM, illustrating the mapping conventions applied across the trehalose headgroup and acyl chains; see **Figures S1-S8** for other lipids. Note that the same lipid color code is used in other similar figures.

### Analysis

For consistency in the analysis, the AA trajectories were mapped onto CG representations using MDAnalysis^40,41^ based on the same CG mapping scheme used in the simulations. All AA-CG comparisons in this study were therefore performed using CG-mapped trajectory generated from the AA simulation together with the native CG trajectory. For the SASA calculations, however, we used the original AA coordinates to prevent the loss of geometry of the surface. Specifically, the *gmx sasa*^*42*^ tool was used to calculate the SASA following the procedure described by Grunewald et al^43^. The van der Waals radii assigned to the MARTINI beads were 2.64 Å for regular beads, 2.30 Å for small beads, and 1.91 Å for tiny beads. For the atomistic simulations, the van der Waals radii were taken from the parameters reported by Rowland and Taylor^44^. A probe radius of 1.91 Å was used for the SASA calculation for both AA and CG simulations.

Pseudo-order parameters and z-dependent pseudo-order parameters were calculated to analyze the phase behavior and lipid acyl chain order (see Supporting Information **S1. Analysis Methods** for details). The mean square displacement (MSD) of each lipid type was calculated using *gmx msd* with a time interval of 1 ns and a time-origin spacing of 1 ns. The diffusion coefficient was obtained by identifying a linear regime of the MSD curve and performing a linear fit to that region. For the AA symmetric α-MA simulations, the diffusion coefficient was taken from Brown et al^15^.

The asymmetric membrane thickness was calculated as the difference between the mean z-coordinates of the lower leaflet’s α-MA headgroups and selected upper leaflet headgroups, excluding PAT and PDIM to avoid positional bias (see **S1. Analysis Methods** for details). The z-density profiles of various lipid components were generated by calculating the headgroup position of each lipid as the center of geometry of specific reference beads relative to the membrane midplane. These individual z-coordinates were then aggregated across all molecules and simulation frames to construct the final density distributions.

The PDIM-PDIM contacts were quantified by calculating pairwise distances between all PDIM molecules (*N*_PDIM_) in each frame of the trajectory. A contact between two PDIM residues is defined when the minimum-image distance (*r*_*c*_) between any beads of residue *a* and residue *b* (|**r**_*a*_(*t*) − **r**_*b*_(*t*)|) is below a cutoff of 6 Å. Any duplicate pairs were removed ({(*a, b*)|*a* ≠ *b*}) to ensure that each unique residue pair was counted only once per frame. For each trajectory, the number of unique contacts per frame was accumulated and averaged over the total number (*N*_frames_) of analyzed frames. The resulting average was further normalized by *N*_PDIM_, yielding the average number of PDIM–PDIM contacts per lipid (*C*_PDIM_).

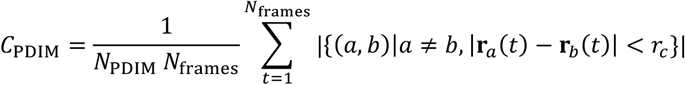

## Results and Discussion

### CG Modeling of *Mycobacterium tuberculosis* Outer Membrane Lipids

We have developed CG models for the major *Mtb* MOM lipids (PDIM, DAT, PAT, SGL, TMM, TDM, and α-MA) based on the MARTINI 3 force field^33,43,45–47^. Each lipid was systematically mapped from its AA representation, accounting for differences in acyl chain length, branching, and functional group placement. Model parameters were refined through an iterative optimization workflow (**Figure 1A**), in which CG simulations were repeatedly compared against AA bonded distributions and updated until convergence was achieved. **Figures 1B** and **1C** show the mapping representation of SGL and TMM, demonstrating the AA-to-CG correspondence and visual consistency of the bead layout; see **Figure S1** for other lipids. The CG mapping and bead naming schemes for all lipids are shown in **Figures 1E** and **S2-S8**.

Parameterizing the trehalose group, which forms the headgroup in the trehalose-based glycolipids (DAT, PAT, SGL, TMM, and TDM), required consideration to capture the ring’s inherent cyclic nature and rigidity, as well as the varied number of acyl chains at the different positions on the trehalose ring. To address these structural features, following previous works^43,45,47^, a TC4 bead was used to represent the cyclic nature of the ring, bond constraints were applied, and one dihedral was used to maintain the planar structure of the rings. The bead type selection within trehalose was the same whenever possible and intentionally non-uniform at the branching location to isolate the –COO– functional group in the acyl chain (**Figures 1B** and **S1**). The bond length within trehalose was scaled up by 15% (**Figure 1D** for T1–T2 in **Figure 1E**) to maintain the stability, without affecting its SASA in the bilayer (**Figure S9**)^43^. This scaling slightly improves the bilayer stability; otherwise, the symmetric outer leaflet membrane showed bilayer instability (data not shown). The only dihedral used in the force field was optimized to address the other three dihedrals (not shown in figures) between sugar groups, which resulted in slightly shifted dihedral distributions in SGL between CG and AA simulations (**Figure S5**).

In addition to the trehalose headgroup mapping, the acyl chains of the MOM lipids are diverse. The C1–C1 bond lengths and C1–C1–C1 bond angles were taken from MARTINI 3,^33^ SC2 bead type was used to represent the beads with methyl group in the acyl chains, SC4 bead (in PAT) was used to represent the bead with a double bond and methyl group, and SC3 bead was chosen for the cyclopropane ring in the mycolate containing lipids. The CG bond length, angle, and dihedral distributions are largely in agreement with the AA counterparts (**Figures S2-S8**). Some bond lengths and bond angles with bimodal distributions were fitted to the midpoint of the distribution due to the limitation of the bonded potential in MARTINI 3. When two peaks differed notably in magnitude, we used the larger population for parameter selection to ensure that the dominant conformational state was captured.

Our CG model captures the structural diversity of *Mtb* MOM lipids, including variations in acyl chain composition, branching, and functional groups, while preserving trehalose ring rigidity and bilayer stability. Iterative refinement against AA distributions ensures that key bonded geometries, SASA, and dihedral preferences are well maintained, despite intrinsic CG limitations in representing bimodal distributions in some bonded parameters. These models provide a robust framework for simulating complex mycobacterial membranes with MARTINI 3, supporting future studies of protein-membrane, membrane-drug, and lipid-lipid interactions.

### Agreement of CG and AA Simulations of *Mtb* Outer Membranes

To evaluate the accuracy, transferability, and physical realism of our CG model, we benchmarked its behavior against AA simulations by comparing key membrane biophysical properties including phase behavior, packing order, thickness, and lipid distribution across various symmetric and asymmetric MOMs.

The previous AA symmetric α-MA membrane exhibits a phase transition around 338 K^15^. To characterize the phase behavior of the CG symmetric α-MA membrane, simulations were performed at multiple temperatures using ten replicas per temperature to ensure statistical robustness and to capture variability associated with phase transitions (**Figure 2A**). Pseudo-order parameter analysis of CG α-MA acyl chain (using the beads in **Figure 2B**) reveals that all ten replicas remain highly ordered at 313 K, 323 K, and 333 K, whereas all ten replicas remain fully disordered at 353 K (**Figure 2C**). At intermediate temperatures, heterogeneity emerges. Five out of ten replicas show markedly reduced order at 338 K, indicating early onset of disorder, while six out of ten replicas become disordered at 343 K (**Figure S10**). This coexistence of ordered and disordered states leads to substantially larger error bars in the pseudo-order parameters at both temperatures, reflecting increased replica-to-replica variability during the gradual phase transition rather than abrupt shift. This observation is consistent with AA simulation showing heterogeneity in phase transition at 338 K, while all three replicas are disordered at 343 K (**Figure 2C)**.

**Figure 2.**
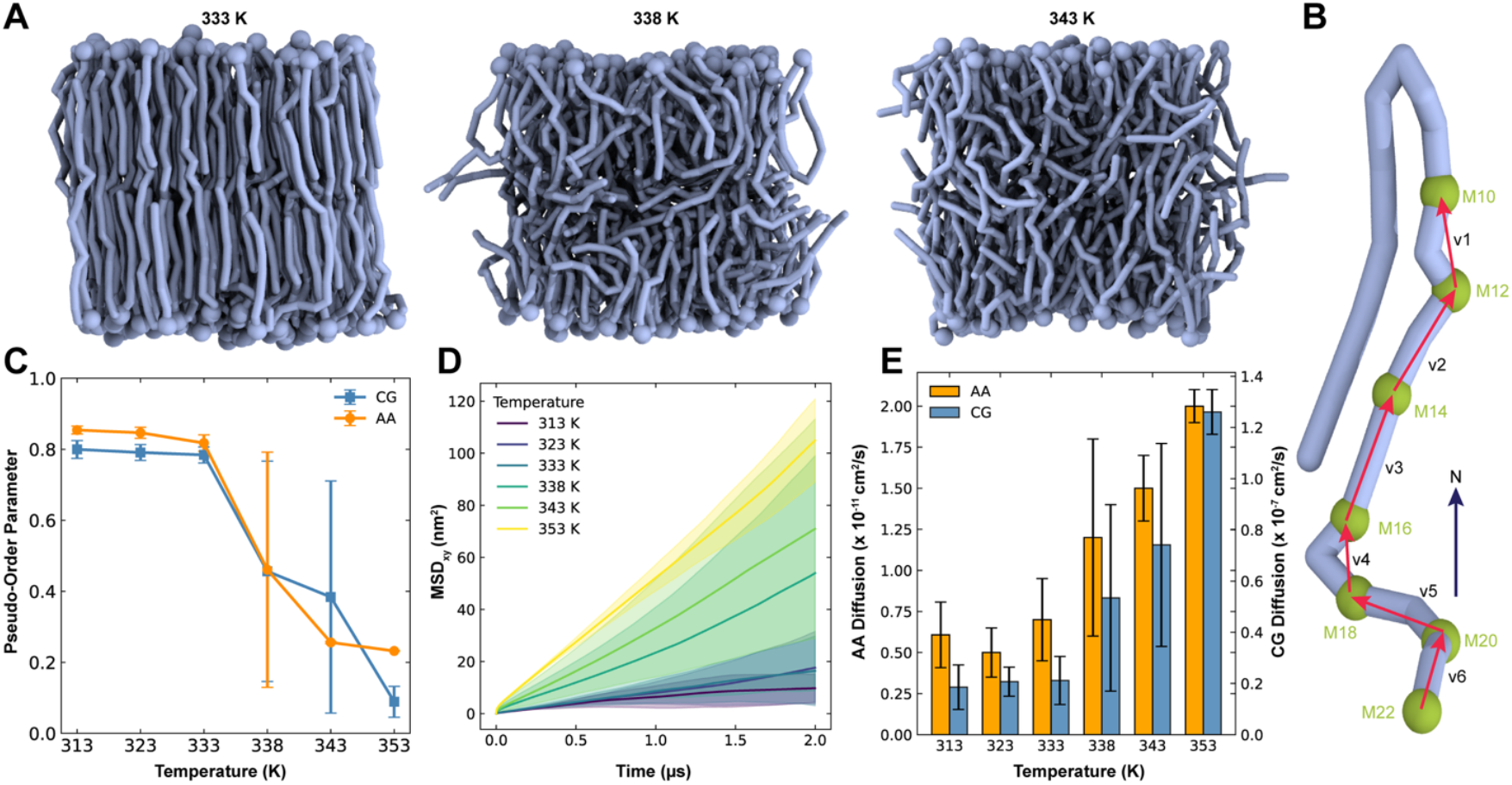
Phase behavior of symmetric α-MA membranes. **A**. Snapshots of equilibrated symmetric α-MA membranes at 333 K, 338 K, and 343 K, illustrating the progression of thermal disorder with increasing temperature. **B**. Schematic showing the selected beads and bond vectors used to compute the α-MA acyl chain pseudo-order parameters. **C**. Temperature-dependent pseudo-order parameter profiles comparing the AA and CG simulations of the symmetric α-MA membrane (three replicas for AA simulations and ten replicas for CG simulations). **D**. Mean-square displacement (MSD) of α-MA in the CG simulations across the examined temperatures, highlighting increased molecular mobility upon heating. **E**. Diffusion coefficients extracted from the AA and CG simulations (0 - 500 ns), demonstrating consistent temperature-dependent trends and the expected systematic overestimation of lipid diffusivity in MARTINI 3 relative to AA simulations.

As expected, lipid diffusion increases with temperature (**Figures 2D, 2E**), with CG diffusion coefficients three orders of magnitude higher than those obtained from the AA simulations. At lower temperatures (i.e., liquid ordered phases), α-MA lipids diffuse slowly and thus their MSD profiles are very noisy and highly variable among the replicas (**Figure S11A**). To explore the system size dependence, additional CG simulations (three replicas) of symmetric α-MA membrane with nine times larger than the initial system were performed (**Figure S12**). While the MSD profiles are still quite variable among the replicas at and before the transition temperature 338 K, diffusion coefficients in the larger CG system are approximately an order magnitude lower than those in the smaller CG system (**Figure S11B**). Although larger systems are often expected to exhibit higher diffusion coefficients due to reduced finite-size effects^48,49^, the opposite trend is observed in our study due to the uncertainty in estimating lipid diffusion from the small system at liquid ordered phases. In other words, the higher diffusion observed in the smaller system at and before the transition temperature should be interpreted cautiously. In contrast, at higher temperatures, i.e., liquid disordered phases, MSD profiles are smoothly linear, and diffusion coefficients are consistent across different CG system sizes (**Figures S11**). Interestingly, α-MA diffusion in liquid disordered phases (at 353 K) is comparable to that of POPC (2.6×10^-7^ cm^2^/s) at 313 K (**Figures 2E, S13**) even with very different acyl chain nature between α-MA and POPC; n.b., POPC CG diffusion coefficient is approximately fourfold higher than the AA one (6.3×10^-8^ cm^2^/s)^50^. Together, these results demonstrate that, while MARTINI 3 overestimates absolute diffusion coefficients of α-MA compared to AA simulations, likely due to long acyl chains of α-MA and smoother energy landscape inherent to coarse-graining, the model nevertheless captures the qualitative trend of increasing diffusion with membrane disorder and reproduces consistent phase-dependent behavior across the system sizes (**Figures 2** and **S12**).

Next, a detailed comparison between the asymmetric MOM AA and CG simulations is performed. The lipid headgroup density profiles from the CG simulations broadly reproduce those from the AA simulations for most lipid species (**Figure 3A**, see also **Figure S14** for three individual CG replicas). While a subset of PAT headgroups is observed near the membrane midplane (z = 0) in the AA simulations, this feature is not evident in the CG system; however, in larger CG systems, a few PAT molecules similarly migrate toward the bilayer center (see next subsection and **Figure 4D**). The PDIM headgroup consistently appears near the membrane center in both AA and CG simulations, likely due to its lack of strong polar groups.

**Figure 3.**
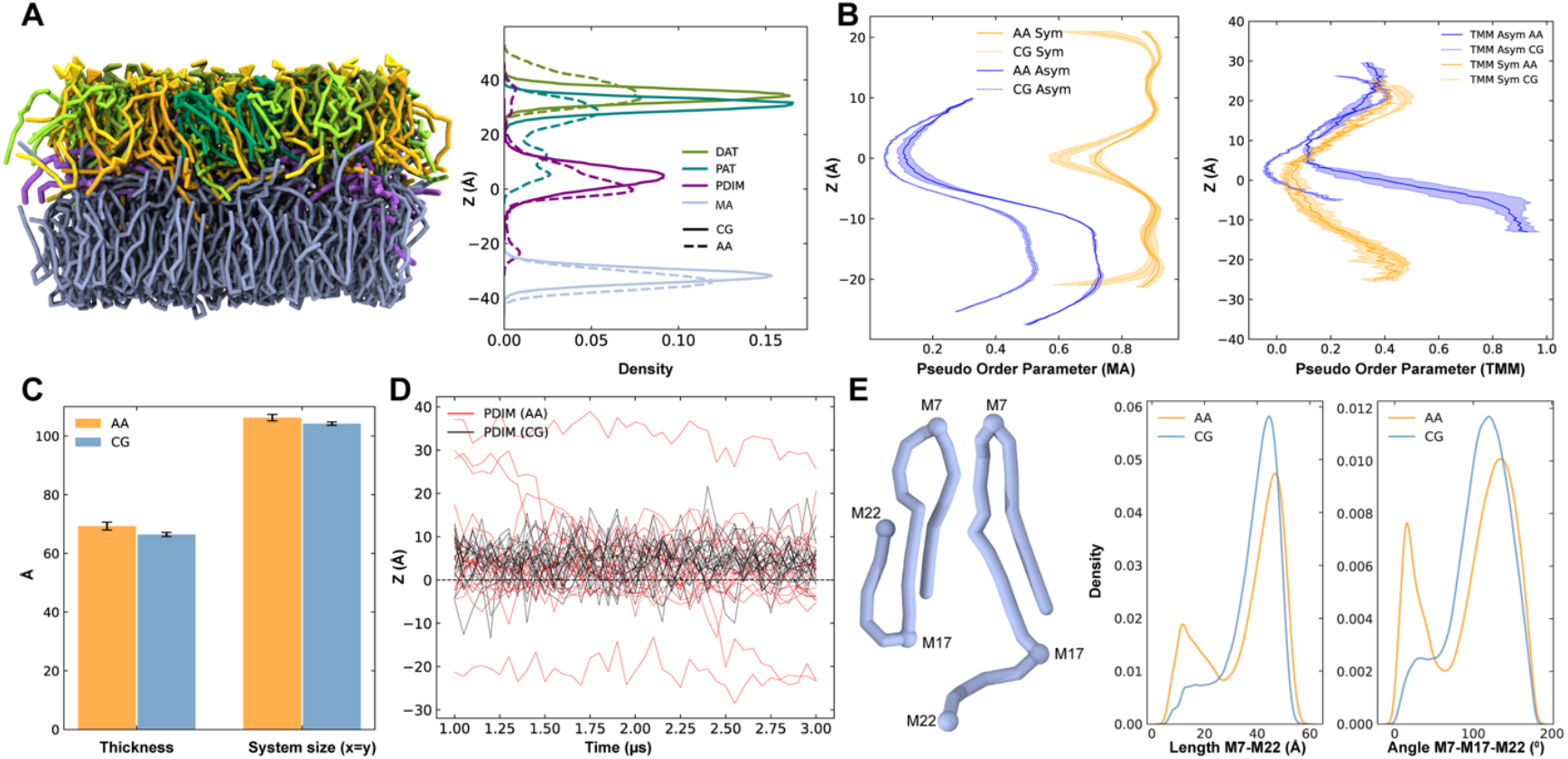
Comparison of membrane biophysical properties between the asymmetric MOM AA and CG simulations. **A**. Equilibrated asymmetric MOM CG structure with the lipid headgroup z-density profiles for DAT (olive), PAT (teal), PDIM (purple), and α-MA (light blue): dashed lines for CG and solid lines for AA. **B**. Z-dependent pseudo-order parameter comparison of α-MA between AA and CG simulations for the symmetric α-MA membrane and the asymmetric MOM, highlighting the reduction in α-MA acyl chain order in the asymmetric system (left). Z-dependent pseudo-order parameter comparison for TMM acyl chain between symmetric outer-leaflet membrane and asymmetric membrane is also shown on the right. **C**. Bilayer thickness comparison between AA and CG simulations. **D**. Timeseries of PDIM headgroup positions, demonstrating PDIM migration toward the bilayer center (z = 0): black for CG and red for AA. **E**. Bead selection for elongated and folded α-MA conformations (left), and distributions of the head– tail distance (M7–M22) and the head–cyclopropane–tail angle (M7–M17–M22); see **Figure S8** for α-MA CG mapping.

**Figure 4.**
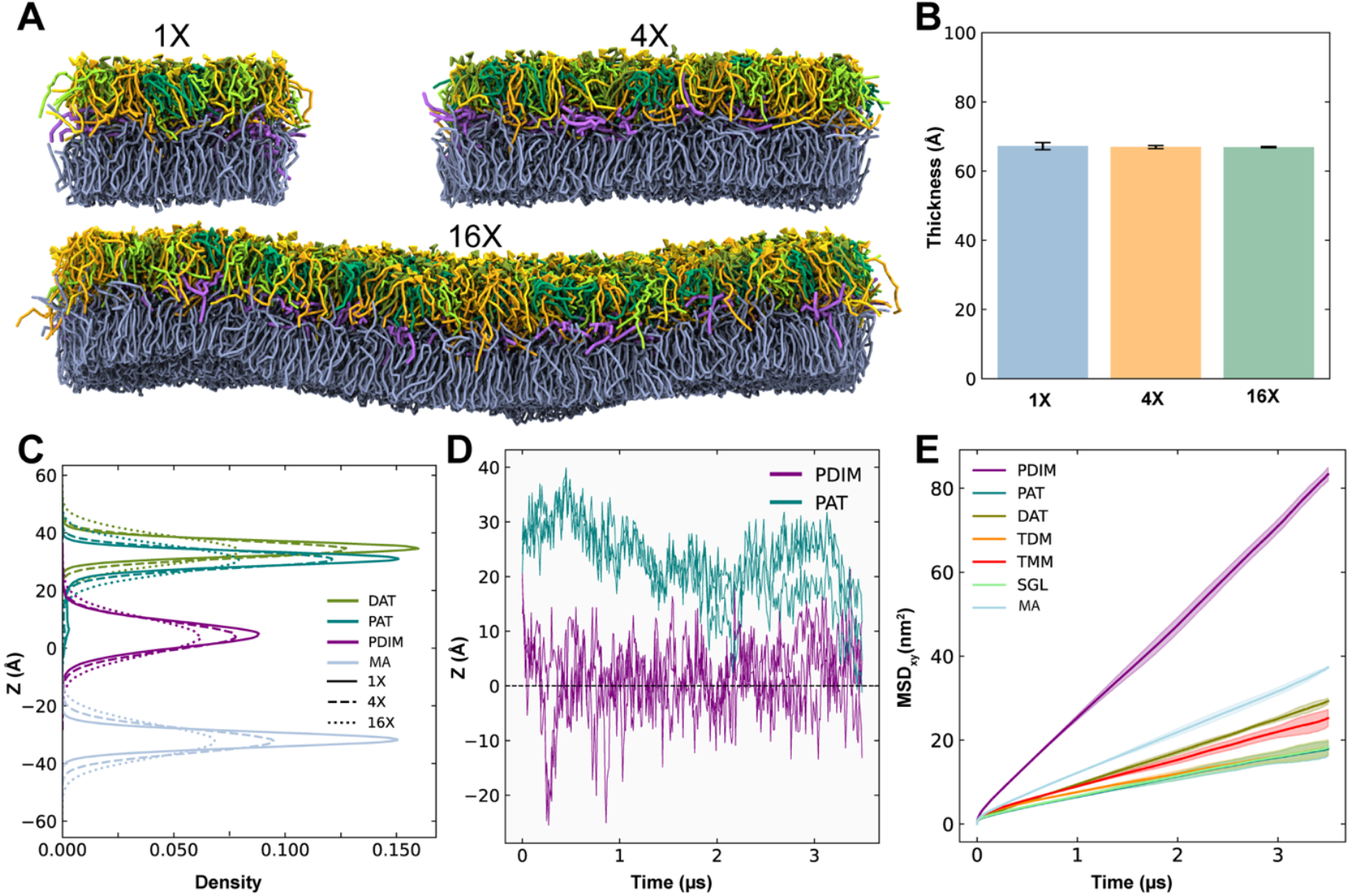
Scalability and structural properties of the asymmetric MOM CG model. **A**. Representative snapshots of equilibrated CG membrane systems simulated at three different scales: 1X (∼120 Å x 120 Å), 4X (∼240 Å x 240 Å), and 16X (∼480 Å x 480 Å) in the xy plan (i.e., the membrane surface area). **B**. Comparison of the average membrane thickness across the three system sizes, showing consistent thickness regardless of scale. **C**. Z-density profiles of lipid headgroups along the membrane normal. The profiles for the 1X (solid lines), 4X (dashed lines), and 16X (dotted lines) systems are overlaid, demonstrating that structural organization and peak positions remain consistent across system sizes. **D**. Timeseries of the z-coordinate for representative PDIM and PAT molecules, highlighting the characteristic migration of PAT and PDIM toward the membrane center (z=0) in 16X system. **E**. MSDs calculated for each lipid type in 16X system.

Comparison of acyl chain pseudo-order parameters as a function of z-position (see **S1. Analysis Methods** for details) reveals that CG simulations exhibit slightly lower ordering than their AA counterparts in both symmetric α-MA and asymmetric MOM (**Figure 3B**). Notably, the α-MA acyl chains in the asymmetric membrane display a marked reduction in order relative to those in the symmetric α-MA bilayer, recapitulating the induced disorder at the inner leaflet observed in the AA simulation, i.e., the effect of outer leaflet lipids in disrupting the order of inner leaflet α-MA (**Figure 3B**, left). In contrast, the pseudo-order parameter of TMM acyl chains in the upper leaflet remains similar between the symmetric and asymmetric systems (**Figure 3B**, right).

The overall bilayer thickness in the CG systems is slightly underestimated compared to AA simulations, through the deviation is minor and the thickness remains comparable to experimental measurements (∼70 - 80 Å)^51,52^ (**Figure 3C**). Timeseries analysis of PDIM headgroup positions reveals that all PDIM molecules migrate toward the bilayer center in the CG simulations (**Figure 3D**, see also **Figure S15** for three individual CG replicas). The AA trajectories also indicate a gradual inward migration of PDIMs, suggesting that complete relocation to the midplane can occur over longer timescales.

Additionally, the AA simulations, which began with elongated, semi-folded, and fully folded α-MA confirmations (2:1:1), showed a progressive shift toward elongated states over time^15^. As shown in **Figure 3E**, the head-to-tail distance and head-cyclopropane-tail angle distributions reveal a broader dominant peak and a smaller secondary peak in the AA simulations, whereas the secondary peak is substantially reduced in the CG simulations. This result indicates that the CG model is more biased toward the elongated confirmation, and the elongated confirmation could be more populated in longer AA simulations. Neighbor count analysis of outer leaflet lipids based on lipid centers of geometry does not indicate any preferential aggregation patterns in both AA and CG systems (**Figure S16**).

Taken together, our CG models of *Mtb* MOM lipids reproduce the essential thermodynamic and structural features of the corresponding AA systems. In the symmetric α-MA bilayers, the CG model captures the heterogeneous phase transition near 338 K and preserves the qualitative temperature dependence of lipid mobility, despite the overestimation of the absolute diffusion compared to AA simulations. In the asymmetric MOM systems, CG simulations recapitulate leaflet-dependent ordering, lipid density distributions, and bilayer thickness observed in the AA simulations, including the outer-leaflet-induced disordering of inner leaflet α-MA. Although PDIM migration and α-MA conformational preferences are enhanced in CG, these differences primarily reflect accelerated dynamics and a smoother free-energy landscape rather than qualitative structural artifacts. Collectively, the models provide a physically realistic and computationally efficient framework for studying the organization and dynamics of MOMs, while maintaining consistency with key qualitative features observed in AA simulations.

### Modulation of PDIM Biophysical Properties by Membrane Fluidity and Composition

Both asymmetric MOM AA and CG simulations show PDIM frequently migrated toward the membrane center (**Figures 3A, 3D** and **4C, 4D**), and prior literature reported PDIM aggregation in POPC bilayers^25^. Interestingly, while PDIM moves to the membrane center, aggregation is not observed in the asymmetric MOM (**Figure S16**), suggesting that PDIM behavior strongly depends on membrane composition. To better understand how PDIM molecules diffuse toward and aggregate in the membrane center, we next investigate their behavior across multiple membrane environments including POPC, PEPC, PSPC, MOM inner leaflet symmetric, MOM outer leaflet symmetric, and MOM asymmetric systems; see **Figure S17** for POPC, PEPC, and PSPC structures. The goal is to determine how differences in membrane fluidity, governed by chain saturation, length, leaflet asymmetry, and temperature, influence PDIM positioning, dynamics, and interactions. POPC, PEPC, and PSPC are chosen to systematically vary membrane fluidity: POPC (16:0, 18:1) contains one unsaturation, PSPC (16:0, 18:0) is fully saturated, and PEPC (18:0, 22:1) has a slightly longer chain than POPC while retaining unsaturation. In addition, symmetric versus asymmetric MOM bilayers mimic the MOM heterogeneity. For these environments, we used CG simulations with larger system sizes and longer timescale that could not be easily simulated using the AA model.

Before investigating these larger systems, we first performed scaling tests on the asymmetric MOM bilayer to verify that our CG model maintains consistent structural and dynamic properties across three different system sizes: ∼120 Å x 120 Å (1X), ∼240 Å x 240 Å (4X), and ∼480 Å x 480 Å (16X) in the xy plane (**Figure 4A**). Across all scales, the membrane thickness remains consistent (**Figure 4B**), and the z-density profiles of lipid headgroups show similar peak positions (**Figure 4C**, see also **Figure S18** for three individual replicas). Although larger systems exhibit slightly broader headgroup distributions, likely due to increased membrane undulations, the overall structural properties remain unchanged. The characteristic movement of PAT toward the membrane center, previously observed in the AA simulations, also occurs in large systems (**Figure 4D**, see also **Figure S19** for three individual replicas in 16X system), though at low frequency. Diffusion behavior is also consistent across scales (**Figures 4E** and **S20**), with PDIM displaying the highest mobility, followed by α-MA, DAT, TMM, TDM, and finally PAT and SGL, indicating that larger or more branched lipids diffuse more slowly. These results confirm that our CG membrane model is scalable, so subsequent simulations of PDIM in various membrane bilayers were therefore performed using the 4X membrane.

To explore how acyl chain saturation and length influence PDIM’s location, dynamics, and clustering, CG simulations were carried out in a series of membranes: POPC, PEPC, PSPC, α-MA-only inner leaflet symmetric, an outer leaflet symmetric membrane, and the full asymmetric MOM (**Figure 5A**). Because these membranes exhibit different phase states, their simulations were conducted at three different temperatures (**Table S2**): 253 K (lower than experimental phase transition for POPC^53^), 313 K (physiological), and 353 K (liquid phase for α-MA). At 253 K, PDIM headgroups in POPC are frequently located near the bilayer center, but increasing chain length (PEPC) or saturation (PSPC) restricts this motion, keeping PDIM headgroups near the membrane-water interface and PDIM tails buried at the membrane center (**Figures 5A, 5B**, see also **Figure S21** for three individual replicas). At 313 K, where POPC and PEPC are fluid, PDIM headgroups are largely localized to the membrane center, whereas in more ordered membranes (PSPC and α-MA), most headgroups remain at the surface, though a secondary density peak emerges in the center for PSPC. At 353 K, where all membranes are fully fluid, PDIM headgroups are consistently migrated to the membrane center, demonstrating that PDIM z-position is strongly governed by membrane fluidity. This fluidity-dependent behavior of hydrophobic lipids is reminiscent with other neutral lipid systems such as triacylglycerols and sterol esters, which are known to accumulate in the hydrophobic core of the membranes and, at sufficient concentrations, phase-separate to form lipid lens and lipid droplets in both model bilayers and cellular systems^54– 58^.

**Figure 5.**
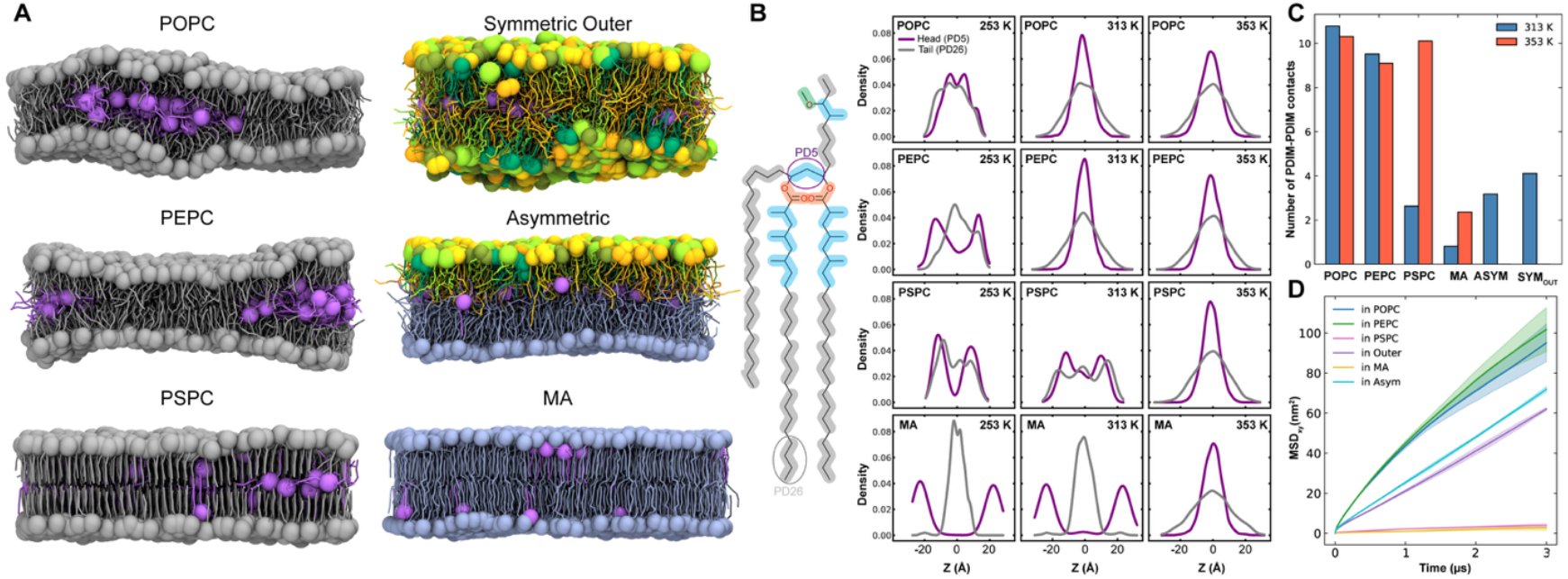
Influence of membrane fluidity and lipid environment on PDIM behavior. **A**. Representative snapshots of PDIM (purple) embedded in various equilibrated lipid bilayers at 313 K: symmetric POPC, PEPC, and PSPC membranes; symmetric outer leaflet and inner leaflet MOM models; and a full asymmetric MOM model. Each lipid headgroup is represented by sphere. **Figure S23** shows both top and side views of each system at three different temperatures. **B**. Z-density profiles of PDIM headgroup bead (PD5, purple) compared to the tail one (PD26, grey) at varying temperatures (253 K, 313 K, and 353 K). The profiles show a shift in PDIM localization from the surface to the membrane center as temperature (and thus fluidity) increases. **C**. Quantification of PDIM-PDIM contacts (within 6 Å) at 313 K and 353 K, showing that aggregation is generally higher in fluid membranes. **D**. MSDs of PDIM in various lipid environment at 313 K, confirming that PDIM diffuses fastest in fluid membranes (POPC and PEPC) and significantly slower in more ordered environments (PSPC and α-MA).

To determine whether membrane fluidity also affects lateral organization, PDIM-PDIM contacts are quantified at 313 K and 353 K (**Figure 5C**, see also **Figure S22** for three individual replicas). At 313 K, strong PDIM aggregation is observed in liquid disordered membranes, with POPC showing ∼10 contacts and PEPC slightly fewer. α-MA exhibits the lowest aggregation, while the asymmetric MOM and outer-leaflet symmetric membranes show higher aggregation than PSPC. These findings indicate that membrane fluidity influences both PDIM z-movement and lateral clustering. At 353 K, substantial PDIM aggregation occurs in PSPC, nearly equal to POPC, whereas aggregation remains low in α-MA membranes even with PDIM in the center. Notably, PEPC shows slightly lower aggregation than PSPC at 353 K, a trend that is highly reproducible across replicas (**Figures S22** and **S23**). Together, this observation suggests that PDIM aggregation is modulated not only by membrane fluidity but possibly also by the hydrophobic acyl chain length of host lipids. Diffusion analyses further support this conclusion that PDIM diffuses fastest in POPC and PEPC membranes and significantly slower in PSPC and α-MA membranes, consistent with restricted movement in more ordered environments (**Figure 5D**). Moderate diffusion in the outer-leaflet symmetric and asymmetric membranes suggests that outer leaflet lipids can enhance inner leaflet fluidity by modifying packing order (**Figure 3B**).

Although PDIM is recognized as a key determinant of mycobacterial virulence and antibiotic tolerance^59–64^, the physical principles governing its organization within membrane remains poorly understood. Our result demonstrates that changes in membrane composition directly modulate PDIM behavior. By explicitly linking membrane biophysics to PDIM organization, this study offers a framework for interpreting how variations in lipid environment influence PDIM-associated membrane properties, with potential implications for mycobacterial physiology and virulence-related phenotypes.

## Conclusions

In this study, we have developed and validated a MARTINI 3 CG model of the *Mtb* outer membrane that captures its pronounced lipid asymmetry and compositional complexity. By parameterizing the major MOM lipids against AA simulations and benchmarking key structural and dynamic membrane properties, we have established a CG framework that reliably reproduces membrane thickness, lipid organization, phase behavior, and density distributions across symmetric and asymmetric AA systems. While our CG model exhibits expected acceleration of lipid diffusion inherent to CG force fields, it preserves relative trends in mobility and ordering across lipid environments.

Application of our CG model reveals that the biophysical behavior of PDIM is strongly governed by membrane fluidity and composition. PDIM preferentially migrates toward the bilayer center in fluid membranes, accompanied by enhanced diffusion and lateral aggregation, whereas ordered environments restrict both translocation and clustering. In addition to membrane fluidity, our results indicate that the acyl chain length further modulates PDIM behavior, influencing both its vertical positioning and lateral interactions. Membranes composed of longer or more rigid lipid species such as α-MA impose stronger constraints on PDIM mobility and aggregation, underscoring the coupled roles of membrane order and hydrophobic thickness in regulating lipid organization. These findings highlight the importance of the surrounding lipid matrix in modulating the spatial distribution and collective behavior of virulence-associated lipids in the MOM.

Overall, our CG MOM model provides a scalable and computationally efficient platform for studying mycobacterial membrane organization at mesoscopic length and time scales. This framework enables systematic investigation of protein-lipid and lipid–lipid interactions, membrane remodeling, and drug permeation mechanisms that are inaccessible to AA simulations alone, and thus offers a valuable tool for future studies of mycobacteria membrane biology and therapeutic targeting.

## Acknowledgements

This work is supported by the CNRS-MITI grant “Modélisation du vivant” 2020 (to M.C.), NSF MCB-2111728, and NIH R35 GM153458 (to W.I.).

## Author Contributions

B.A, M.C., and W.I. designed the research. B.A. performed the research and drafted the paper with input from the co-authors. S.L., T.P.B., M.C., and W.I. helped with interpretation of results and revised the manuscript.

## Data Availability

The input, some trajectory, and restart files, as well as some analysis code are freely available in a Zenodo dataset (https://doi.org/10.5281/zenodo.18671377).

## References

(1) Cambau, E.; Drancourt, M. Steps towards the Discovery of Mycobacterium Tuberculosis by Robert Koch, 1882. Clinical Microbiology and Infection 2014, 20 (3), 196–201. 10.1111/1469-0691.12555.

(2) Russell, D. G. Mycobacterium Tuberculosis: Here Today, and Here Tomorrow. Nature Reviews Molecular Cell Biology 2001 2:8 2001, 2 (8), 569–578. 10.1038/35085034.

(3) Global Tuberculosis Report 2025. https://www.who.int/teams/global-programme-on-tuberculosis-and-lung-health/tb-reports/global-tuberculosis-report-2025 (accessed 2026-02-14).

(4) Dheda, K.; Mirzayev, F.; Cirillo, D. M.; Udwadia, Z.; Dooley, K. E.; Chang, K. C.; Omar, S. V.; Reuter, A.; Perumal, T.; Horsburgh, C. R.; Murray, M.; Lange, C. Multidrug-Resistant Tuberculosis. Nature Reviews Disease Primers 2024 10:1 2024, 10 (1), 22-. 10.1038/s41572-024-00504-2.

(5) Seung, K. J.; Keshavjee, S.; Rich, M. L. Multidrug-Resistant Tuberculosis and Extensively Drug-Resistant Tuberculosis. Cold Spring Harb. Perspect. Med. 2015, 5 (9), a017863. 10.1101/CSHPERSPECT.A017863.

(6) Gray, R. M.; Hunt, D. M.; Silva dos Santos, M.; Liu, J.; Agapova, A.; Rodgers, A.; Fearns, A.; Canseco, J. O.; Garza-Garcia, A.; MacRae, J. I.; Gutierrez, M. G.; Lee, R. E.; de Carvalho, L. P. S. Mycobacterium Tuberculosis Overcomes Phosphate Starvation by Extensively Remodelling Its Lipidome with Phosphorus-Free Lipids. Nature Communications 2025 16:1 2025, 16 (1), 11317-. 10.1038/s41467-025-66437-w.

(7) Schami, A.; Islam, M. N.; Belisle, J. T.; Torrelles, J. B. Drug-Resistant Strains of Mycobacterium Tuberculosis: Cell Envelope Profiles and Interactions with the Host. Front. Cell. Infect. Microbiol. 2023, 13, 1274175. 10.3389/FCIMB.2023.1274175.

(8) Garcia-Vilanova, A.; Chan, J.; Torrelles, J. B. Underestimated Manipulative Roles of Mycobacterium Tuberculosis Cell Envelope Glycolipids During Infection. Front. Immunol. 2019, 10. 10.3389/FIMMU.2019.02909.

(9) Batt, S. M.; Minnikin, D. E.; Besra, G. S. The Thick Waxy Coat of Mycobacteria, a Protective Layer against Antibiotics and the Host’s Immune System. Biochemical Journal 2020, 477 (10), 1983–2006. 10.1042/BCJ20200194.

(10) Dulberger, C. L.; Rubin, E. J.; Boutte, C. C. The Mycobacterial Cell Envelope — a Moving Target. Nature Reviews Microbiology 2019 18:1 2019, 18 (1), 47–59. 10.1038/s41579-019-0273-7.

(11) Jackson, M. The Mycobacterial Cell Envelope—Lipids. Cold Spring Harb. Perspect. Med. 2014, 4 (10), a021105. 10.1101/CSHPERSPECT.A021105.

(12) Karakousis, P. C.; Bishai, W. R.; Dorman, S. E. Mycobacterium Tuberculosis Cell Envelope Lipids and the Host Immune Response. Cell. Microbiol. 2004, 6 (2), 105–116. 10.1046/j.1462-5822.2003.00351.x.

(13) Lee, R. E.; Brennan, P. J.; Besra, G. S. Mycobacterium Tuberculosis Cell Envelope. Curr. Top. Microbiol. Immunol. 1996, 215, 1–27. 10.1007/978-3-642-80166-2_1.

(14) Liang, K.; Mathew, L.; Fu, L.; Srivatsav, A.; Duan, J.; Ni, L.; Luo, Z.; Kapoor, S.; Duan, M. Development of Force Field Parameters for the Atomic Simulation of Mycobacterial Membranes. Communications Chemistry 2025 8:1 2025, 8 (1), 345-. 10.1038/s42004-025-01722-9.

(15) Brown, T. P.; Chavent, M.; Im, W. Dynamic Architecture of Mycobacterial Outer Membranes Revealed by All-Atom Simulations. Elife 2025, 14. 10.7554/ELIFE.108644.1.

(16) Brown, T.; Chavent, M. G.; Im, W. Molecular Dynamics Simulation of Mycobacterium Tuberculosis Cell Envelopes: Mycomembranes. Biophys. J. 2024, 123 (3), 507a. 10.1016/j.bpj.2023.11.3063.

(17) Vasyankin, A. V.; Panteleev, S. V.; Steshin, I. S.; Shirokova, E. A.; Rozhkov, A. V.; Livshits, G. D.; Radchenko, E. V.; Ignatov, S. K.; Palyulin, V. A. Temperature-Induced Restructuring of Mycolic Acid Bilayers Modeling the Mycobacterium Tuberculosis Outer Membrane: A Molecular Dynamics Study. Molecules 2024, 29 (3), 696. 10.3390/MOLECULES29030696.

(18) Paes, Y. M.; Scaini, J. L. R.; de Lima, V. R.; Werhli, A. V.; da Silva, P. A.; Bordin, J. R.; Machado, K. dos S. A Molecular Dynamics Study of PIM2 Lipid Bilayer Membranes. J. Mol. Graph. Model. 2026, 144, 109284. 10.1016/J.JMGM.2026.109284.

(19) Kumar, Y.; Basu, S.; Chatterji, D.; Ghosh, A.; Jayaraman, N.; Maiti, P. K. Self-Assembly of Mycolic Acid in Water: Monolayer or Bilayer. Langmuir 2025, 41 (5), 3140–3156. 10.1021/ACS.LANGMUIR.4C03743.

(20) Li, Y.; Acharya, A.; Yang, L.; Liu, J.; Tajkhorshid, E.; Zgurskaya, H. I.; Jackson, M.; Gumbart, J. C. Insights into Substrate Transport and Water Permeation in the Mycobacterial Transporter MmpL3. Biophys. J. 2023, 122 (11), 2342–2352. 10.1016/J.BPJ.2023.03.018.

(21) Adhyapak, P.; Dong, W.; Dasgupta, S.; Dutta, A.; Duan, M.; Kapoor, S. Lipid Clustering in Mycobacterial Cell Envelope Layers Governs Spatially Resolved Solvation Dynamics. Chem. Asian J. 2022, 17 (11), e202200146. 10.1002/ASIA.202200146.

(22) Marrink, S. J.; Corradi, V.; Souza, P. C. T.; Ingólfsson, H. I.; Tieleman, D. P.; Sansom, M. S. P. Computational Modeling of Realistic Cell Membranes. Chem. Rev. 2019, 119 (9), 6184–6226. 10.1021/ACS.CHEMREV.8B00460.

(23) Brown, C. M.; Marrink, S. J. Modeling Membranes in Situ. Curr. Opin. Struct. Biol. 2024, 87, 102837. 10.1016/J.SBI.2024.102837.

(24) Brown, C. M.; Westendorp, M. S. S.; Zarmiento-Garcia, R.; Stevens, J. A.; Bruininks, B. M. H.; Rouse, S. L.; Marrink, S. J.; Wassenaar, T. A. An Integrative Modelling Approach to the Mitochondrial Cristae. Communications Biology 2025 8:1 2025, 8 (1), 972-. 10.1038/s42003-025-08381-5.

(25) Augenstreich, J.; Haanappel, E.; Ferré, G.; Czaplicki, G.; Jolibois, F.; Destainville, N.; Guilhot, C.; Milon, A.; Astarie-Dequeker, C.; Chavent, M. The Conical Shape of DIM Lipids Promotes Mycobacterium Tuberculosis Infection of Macrophages. Proc. Natl. Acad. Sci. U. S. A. 2019, 116 (51), 25649–25658. 10.1073/PNAS.1910368116.

(26) Schahl, A.; Réat, V.; Malaga, W.; Birbes, C.; Czaplicki, G.; Jolibois, F.; Yamamoto, E.; Ramos, P.; Milon, A.; Saurel, O.; Atkinson, R. A.; Astarie-Dequeker, C.; Guilhot, C.; Ferré, G.; Chavent, M.; Haanappel, E. How PGL Finds a Sweet Spot in Phospholipid Membranes: A Combined Multiscale MD and NMR Study. Biophys. J. 2025, 125 (2), 457–470. 10.1016/j.bpj.2025.07.040.

(27) Jo, S.; Kim, T.; Iyer, V. G.; Im, W. CHARMM-GUI: A Web-Based Graphical User Interface for CHARMM. J. Comput. Chem. 2008, 29 (11), 1859–1865. 10.1002/JCC.20945.

(28) Wu, E. L.; Cheng, X.; Jo, S.; Rui, H.; Song, K. C.; Dávila-Contreras, E. M.; Qi, Y.; Lee, J.; Monje-Galvan, V.; Venable, R. M.; Klauda, J. B.; Im, W. CHARMM-GUI Membrane Builder toward Realistic Biological Membrane Simulations. J. Comput. Chem. 2014, 35 (27), 1997–2004. 10.1002/JCC.23702.

(29) Lee, J.; Cheng, X.; Jo, S.; MacKerell, A. D.; Klauda, J. B.; Im, W. CHARMM-GUI Input Generator for NAMD, Gromacs, Amber, Openmm, and CHARMM/OpenMM Simulations Using the CHARMM36 Additive Force Field. Biophys. J. 2016, 110 (3), 641a. 10.1016/j.bpj.2015.11.3431.

(30) Lee, J.; Patel, D. S.; Ståhle, J.; Park, S. J.; Kern, N. R.; Kim, S.; Lee, J.; Cheng, X.; Valvano, M. A.; Holst, O.; Knirel, Y. A.; Qi, Y.; Jo, S.; Klauda, J. B.; Widmalm, G.; Im, W. CHARMM-GUI Membrane Builder for Complex Biological Membrane Simulations with Glycolipids and Lipoglycans. J. Chem. Theory Comput. 2019, 15 (1), 775–786. 10.1021/ACS.JCTC.8B01066.

(31) Abraham, M. J.; Murtola, T.; Schulz, R.; Páll, S.; Smith, J. C.; Hess, B.; Lindah, E. GROMACS: High Performance Molecular Simulations through Multi-Level Parallelism from Laptops to Supercomputers. SoftwareX 2015, 1–2, 19–25. 10.1016/J.SOFTX.2015.06.001.

(32) Abraham, M.; Alekseenko, A.; Bergh, C.; Blau, C.; Briand, E.; Doijade, M.; Fleischmann, S.; Gapsys, V.; Garg, G.; Gorelov, S.; Gouaillardet, G.; Gray, A.; Irrgang, M. E.; Jalalypour, F.; Jordan, J.; Junghans, C.; Kanduri, P.; Keller, S.; Kutzner, C.; Lemkul, J. A.; Lundborg, M.; Merz, P.; Miletić, V.; Morozov, D.; Páll, S.; Schulz, R.; Shirts, M.; Shvetsov, A.; Soproni, B.; Spoel, D. van der; Turner, P.; Uphoff, C.; Villa, A.; Wingbermühle, S.; Zhmurov, A.; Bauer, P.; Hess, B.; Lindahl, E. GROMACS 2023.3 Manual. 10.5281/ZENODO.10017699.

(33) Pedersen, K. B.; Ingólfsson, H. I.; Ramirez-Echemendia, D. P.; Borges-Araújo, L.; Andreasen, M. D.; Empereur-mot, C.; Melcr, J.; Ozturk, T. N.; Bennett, W. F. D.; Kjølbye, L. R.; Brasnett, C.; Corradi, V.; Khan, H. M.; Cino, E. A.; Crowley, J.; Kim, H.; Fábián, B.; Borges-Araújo, A. C.; Pavan, G. M.; Launay, G.; Lolicato, F.; Wassenaar, T. A.; Melo, M. N.; Thallmair, S.; Carpenter, T. S.; Monticelli, L.; Tieleman, D. P.; Schiøtt, B.; Souza, P. C. T.; Marrink, S. J. The Martini 3 Lipidome: Expanded and Refined Parameters Improve Lipid Phase Behavior. ACS Cent. Sci. 2025, 11 (9), 1598–1610. 10.1021/ACSCENTSCI.5C00755.

(34) Bussi, G.; Donadio, D.; Parrinello, M. Canonical Sampling through Velocity Rescaling. Journal of Chemical Physics 2007, 126 (1). 10.1063/1.2408420/186581.

(35) Bernetti, M.; Bussi, G. Pressure Control Using Stochastic Cell Rescaling. Journal of Chemical Physics 2020, 153 (11). 10.1063/5.0020514.

(36) Hess, B.; Bekker, H.; Berendsen, H. J. C.; Fraaije, J. G. E. M. LINCS: A Linear Constraint Solver for Molecular Simulations. J Comput Chem 1997, 18, 14631472. 10.1002/(SICI)1096-987X(199709)18:12<1463::AID-JCC4>3.0.CO;2-H.

(37) Hess, B. P-LINCS: A Parallel Linear Constraint Solver for Molecular Simulation. J. Chem. Theory Comput. 2008, 4 (1), 116–122. 10.1021/CT700200B.

(38) Humphrey, W.; Dalke, A.; Schulten, K. VMD: Visual Molecular Dynamics. J. Mol. Graph. 1996, 14 (1), 33–38. 10.1016/0263-7855(96)00018-5.

(39) Brasnett, C.; Marrink, S. J. MartiniGlass: A Tool for Enabling Visualization of Coarse-Grained Martini Topologies. J. Chem. Inf. Model. 2025, 65 (7), 3137–3141. 10.1021/ACS.JCIM.4C02277.

(40) Michaud-Agrawal, N.; Denning, E. J.; Woolf, T. B.; Beckstein, O. MDAnalysis: A Toolkit for the Analysis of Molecular Dynamics Simulations. J. Comput. Chem. 2011, 32 (10), 2319–2327. 10.1002/JCC.21787.

(41) Gowers, R. J.; Linke, M.; Barnoud, J.; Reddy, T. J. E.; Melo, M. N.; Seyler, S. L.; Domański, J.; Dotson, D. L.; Buchoux, S.; Kenney, I. M.; Beckstein, O. MDAnalysis: A Python Package for the Rapid Analysis of Molecular Dynamics Simulations. SciPy 2016 2016, 98–105. 10.25080/MAJORA-629E541A-00E.

(42) Eisenhaber, F.; Lijnzaad, P.; Argos, P.; Sander, C.; Scharf, M. The Double Cubic Lattice Method: Efficient Approaches to Numerical Integration of Surface Area and Volume and to Dot Surface Contouring of Molecular Assemblies. J. Comput. Chem. 1995, 16 (3), 273–284. 10.1002/JCC.540160303.

(43) Grünewald, F.; Punt, M. H.; Jefferys, E. E.; Vainikka, P. A.; König, M.; Virtanen, V.; Meyer, T. A.; Pezeshkian, W.; Gormley, A. J.; Karonen, M.; Sansom, M. S. P.; Souza, P. C. T.; Marrink, S. J. Martini 3 Coarse-Grained Force Field for Carbohydrates. J. Chem. Theory Comput. 2022, 18 (12), 7555–7569. 10.1021/ACS.JCTC.2C00757.

(44) Rowland, R. S.; Taylor, R. Intermolecular Nonbonded Contact Distances in Organic Crystal Structures: Comparison with Distances Expected from van Der Waals Radii. 1996. 10.1021/JP953141.

(45) Alessandri, R.; Barnoud, J.; Gertsen, A. S.; Patmanidis, I.; De Vries, A. H.; Souza, P. C. T.; Marrink, S. J.; Alessandri [+], R.; Barnoud, J.; Patmanidis, I.; De Vries, A. H.; Souza, P. C. T.; Marrink, S. J.; Gertsen, A. S. Martini 3 Coarse-Grained Force Field: Small Molecules. Adv. Theory Simul. 2022, 5 (1), 2100391. 10.1002/ADTS.202100391.

(46) Souza, P. C. T.; Alessandri, R.; Barnoud, J.; Thallmair, S.; Faustino, I.; Grünewald, F.; Patmanidis, I.; Abdizadeh, H.; Bruininks, B. M. H.; Wassenaar, T. A.; Kroon, P. C.; Melcr, J.; Nieto, V.; Corradi, V.; Khan, H. M.; Domański, J.; Javanainen, M.; Martinez-Seara, H.; Reuter, N.; Best, R. B.; Vattulainen, I.; Monticelli, L.; Periole, X.; Tieleman, D. P.; de Vries, H.; Marrink, S. J. Martini 3: A General Purpose Force Field for Coarse-Grained Molecular Dynamics. Nature Methods 2021 18:4 2021, 18 (4), 382–388. 10.1038/s41592-021-01098-3.

(47) Brandner, A. F.; Smith, I. P. S.; Marrink, S. J.; Souza, P. C. T.; Khalid, S. Systematic Approach to Parametrization of Disaccharides for the Martini 3 Coarse-Grained Force Field. J. Chem. Inf. Model. 2025, 65 (3), 1537–1548. 10.1021/ACS.JCIM.4C01874.

(48) Jamali, S. H.; Wolff, L.; Becker, T. M.; Bardow, A.; Vlugt, T. J. H.; Moultos, O. A. Finite-Size Effects of Binary Mutual Diffusion Coefficients from Molecular Dynamics. J. Chem. Theory Comput. 2018, 14 (5), 2667–2677. 10.1021/ACS.JCTC.8B00170/SUPPL_FILE/CT8B00170_SI_003.ZIP.

(49) Vögele, M.; Hummer, G. Divergent Diffusion Coefficients in Simulations of Fluids and Lipid Membranes. Journal of Physical Chemistry B 2016, 120 (33), 8722–8732. 10.1021/ACS.JPCB.6B05102.

(50) Wang, Y.; Markwick, P. R. L.; De Oliveira, C. A. F.; McCammon, J. A. Enhanced Lipid Diffusion and Mixing in Accelerated Molecular Dynamics. J. Chem. Theory Comput. 2011, 7 (10), 3199. 10.1021/CT200430C.

(51) Hoffmann, C.; Leis, A.; Niederweis, M.; Plitzko, J. M.; Engelhardt, H. Disclosure of the Mycobacterial Outer Membrane: Cryo-Electron Tomography and Vitreous Sections Reveal the Lipid Bilayer Structure. Proc. Natl. Acad. Sci. U. S. A. 2008, 105 (10), 3963. 10.1073/PNAS.0709530105.

(52) Zuber, B.; Chami, M.; Houssin, C.; Dubochet, J.; Griffiths, G.; Daffé, M. Direct Visualization of the Outer Membrane of Mycobacteria and Corynebacteria in Their Native State. J. Bacteriol. 2008, 190 (16), 5672–5680. 10.1128/JB.01919-07.

(53) Phase Transition Temperatures for Glycerophospholipids | Avanti Research. https://www.avantiresearch.com/en-gb/support-hub/physical-properties/phase-transition-temps (accessed 2026-01-02).

(54) Olzmann, J. A.; Carvalho, P. Dynamics and Functions of Lipid Droplets. Nature Reviews Molecular Cell Biology 2018 20:3 2018, 20 (3), 137–155. 10.1038/s41580-018-0085-z.

(55) Shpilka, T.; Welter, E.; Borovsky, N.; Amar, N.; Mari, M.; Reggiori, F.; Elazar, Z. Lipid Droplets and Their Component Triglycerides and Steryl Esters Regulate Autophagosome Biogenesis. EMBO J. 2015, 34 (16), 2117–2131. 10.15252/EMBJ.201490315.

(56) Kuerschner, L.; Moessinger, C.; Thiele, C. Imaging of Lipid Biosynthesis: How a Neutral Lipid Enters Lipid Droplets. Traffic 2008, 9 (3), 338–352. 10.1111/J.1600-0854.2007.00689.X.

(57) Park, S.; Choi, Y. K.; Kim, S.; Lee, J.; Im, W. CHARMM-GUI Membrane Builder for Lipid Nanoparticles with Ionizable Cationic Lipids and PEGylated Lipids. J. Chem. Inf. Model. 2021, 61 (10), 5192–5202. 10.1021/ACS.JCIM.1C00770.

(58) Gee, S.; Glover, K. J.; Wittenberg, N. J.; Im, W. CHARMM-GUI Membrane Builder for Lipid Droplet Modeling and Simulation. Chempluschem 2024, 89 (8), e202400013. 10.1002/CPLU.202400013.

(59) Warner, D. F.; Barczak, A. K.; Gutierrez, M. G.; Mizrahi, V. Mycobacterium Tuberculosis Biology, Pathogenicity, and Interaction with the Host. Nat. Rev. Microbiol. 2025, 23 (12), 788. 10.1038/S41579-025-01201-X.

(60) Rens, C.; Chao, J. D.; Sexton, D. L.; Tocheva, E. I.; Av-Gay, Y. Roles for Phthiocerol Dimycocerosate Lipids in Mycobacterium Tuberculosis Pathogenesis. Microbiology (United Kingdom) 2021, 167 (3), 001042. 10.1099/MIC.0.001042.

(61) Mittal, E.; Philips, J. A. The Mycobacterium Tuberculosis Lipid, PDIM, Inhibits the NADPH Oxidase and Autophagy. Autophagy 2025, 21 (3), 684–685. 10.1080/15548627.2024.2439928.

(62) Block, A. M.; Namugenyi, S. B.; Palani, N. P.; Brokaw, A. M.; Zhang, L.; Beckman, K. B.; Tischler, A. D. Mycobacterium Tuberculosis Requires the Outer Membrane Lipid Phthiocerol Dimycocerosate for Starvation-Induced Antibiotic Tolerance. mSystems 2023, 8 (1). 10.1128/msystems.00699-22.

(63) Mittal, E.; Prasad, G. V. R. K.; Upadhyay, S.; Sadadiwala, J.; Olive, A. J.; Yang, G.; Sassetti, C. M.; Philips, J. A. Mycobacterium Tuberculosis Virulence Lipid PDIM Inhibits Autophagy in Mice. Nature Microbiology 2024 9:11 2024, 9 (11), 2970–2984. 10.1038/s41564-024-01797-5.

(64) Astarie-Dequeker, C.; Le Guyader, L.; Malaga, W.; Seaphanh, F. K.; Chalut, C.; Lopez, A.; Guilhot, C. Phthiocerol Dimycocerosates of M. Tuberculosis Participate in Macrophage Invasion by Inducing Changes in the Organization of Plasma Membrane Lipids. PLoS Pathog. 2009, 5 (2), e1000289. 10.1371/JOURNAL.PPAT.1000289.

