## Supporting Information for "Coarse-Grained Simulations of Mycobacterial Outer Membranes Reveal Fluidity-Dependent PDIM Redistribution Across Different Lipid Environments"

### S1. Analysis Methods

#### Pseudo-order Parameter

The beads used for this calculation are shown in **Fig 2B**. For each bond between adjacent tail beads, the bond vector was computed and normalized, and the pseudo-order parameter was evaluated using the second Legendre polynomial:

$$S_i = \frac{\sum_{t=1}^{N_{\text{frames}}} \sum_{t=1}^{N_{\text{bonds}}} \frac{1}{2}\left(3\left(\hat{v}_{i,t} \cdot \mathbf{n}\right)^2 - 1\right)}{N_{\text{frames}} N_{\text{bonds}}}$$

where the normalized bond vector ($\hat{v}_{i,t}$) and membrane normal ($\mathbf{n}$) are taken as:

$$\hat{v}_{i,t} = \frac{\mathbf{r}_{i+1,t} - \mathbf{r}_{i,t}}{||\mathbf{r}_{i+1,t} - \mathbf{r}_{i,t}||} \quad \text{and} \quad \mathbf{n} = (0,0,1)$$

#### Z-Resolved Pseudo-Order Parameter Calculation

To compute the z-dependent pseudo-order parameter profile *S(z),* the same tail beads were used for mycolates and corresponding beads to mycolates were used for TMM. However, instead of global averaging, the order parameter was evaluated within spatial bins along the membrane normal (z). The midpoint of each bond was assigned to one of 40 equally spaced z-bins and then mean order parameter within each bin was computed:

$$S_i(z_k) = \frac{1}{N_k} \sum_{t=1}^{N_{\text{frames}}} \sum_{i \in \text{bin}(z_k,t)} \frac{1}{2}\left(3\left(\hat{v}_{i,t} \cdot \mathbf{n}\right)^2 - 1\right)$$

#### Thickness Calculation

The z-coordinates of the virtual-site headgroup beads were used as the reference points for the upper leaflet for thickness calculation. Specifically, the headgroup virtual sites of trehalose for TMM (T4 and T8), TDM (TD4 and TD8), and DAT (D4 and D8), as well as the charged bead of SGL (S1) were included; see these bead types in **Figures S1-S8**. PAT and PDIM were excluded because these lipids occasionally migrated toward the bilayer midplane, which would bias the leaflet position. For the lower leaflet, the charged headgroup bead of mycolates (M7) was used as the reference. The bilayer thickness for each frame was computed as the absolute difference between the mean z-coordinates of the upper and lower leaflet reference beads.

**Table S1.** Lipid compositions of the simulated membrane systems.

| System | Resolution | Leaflet | Composition (no of lipids) | | | | | | |
|---|---|---|---|---|---|---|---|---|---|
| | | | DAT | PAT | PDIM | SGL | TDM | TMM | α-MA |
| Symmetric Outer leaflet | AA/CG | Upper | 20 | 20 | 20 | 20 | 20 | 20 | 0 |
| | | Lower | 20 | 20 | 20 | 20 | 20 | 20 | 0 |
| Symmetric Inner leaflet | AA/CG | Upper | 0 | 0 | 0 | 0 | 0 | 0 | 60 |
| | | Lower | 0 | 0 | 0 | 0 | 0 | 0 | 60 |
| Symmetric Inner leaflet (9X) | CG | Upper | 0 | 0 | 0 | 0 | 0 | 0 | 540 |
| | | Lower | 0 | 0 | 0 | 0 | 0 | 0 | 540 |
| Asymmetric Outer membrane (1X) | AA/CG | Upper | 20 | 20 | 20 | 20 | 20 | 20 | 0 |
| | | Lower | 0 | 0 | 0 | 0 | 0 | 0 | 188 |
| Asymmetric Outer membrane (4X) | CG | Upper | 80 | 80 | 80 | 80 | 80 | 80 | 0 |
| | | Lower | 0 | 0 | 0 | 0 | 0 | 0 | 752 |
| Asymmetric Outer membrane (16X) | CG | Upper | 320 | 320 | 320 | 320 | 320 | 320 | 0 |
| | | Lower | 0 | 0 | 0 | 0 | 0 | 0 | 3008 |
| PDIM in Symmetric Outer leaflet | CG | Upper | 80 | 80 | 40 | 80 | 80 | 80 | 0 |
| | | Lower | 80 | 80 | 40 | 80 | 80 | 80 | 0 |
| PDIM in Asymmetric Outer membrane | CG | Upper | 80 | 80 | 80 | 80 | 80 | 80 | 0 |
| | | Lower | 0 | 0 | 0 | 0 | 0 | 0 | 752 |
| PDIM in MA | CG | Upper | 0 | 0 | 40 | 0 | 0 | 0 | 760 |
| | | Lower | 0 | 0 | 40 | 0 | 0 | 0 | 760 |

| | | | PDIM | POPC | PEPC | PSPC |
|---|---|---|---|---|---|---|
| POPC only | CG | Upper | 0 | 100 | 0 | 0 |
| | | Lower | 0 | 100 | 0 | 0 |
| PDIM in POPC | CG | Upper | 40 | 760 | 0 | 0 |
| | | Lower | 40 | 760 | 0 | 0 |
| PDIM in PEPC | CG | Upper | 40 | 0 | 760 | 0 |
| | | Lower | 40 | 0 | 760 | 0 |
| PDIM in PSPC | CG | Upper | 40 | 0 | 0 | 760 |
| | | Lower | 40 | 0 | 0 | 760 |

**Table S2.** Simulation system information.

| System | # total atoms | Initial system size in X Y Z (Å) | Simulation time (μs) | # CG replicas[a] | Temperature |
|---|---|---|---|---|---|
| Symmetric Outer leaflet | 23079 | 133 133 137 | 3 | 3 | 313 K |
| Symmetric Inner leaflet | 4714 | 77 77 106 | 3 | 10 | 313 K, 323 K, 333 K, 338 K, 353 K |
| Symmetric Inner leaflet (9X) | 40716 | 217 217 100 | 3 | 3 | 313 K, 323 K, 333 K, 338 K, 353 K |
| Asymmetric Outer membrane (1X) | 15961 | 119 119 114 | 10 | 3 | 313 K |
| Asymmetric Outer membrane (4X) | 63844 | 239 239 114 | 10 | 3 | 313 K |
| Asymmetric Outer membrane (16X) | 255376 | 479 479 479 | 3.5 | 3 | 313 K |
| PDIM in Symmetric Outer leaflet | 75557 | 235 235 131 | 3 | 3 | 313 K |
| PDIM in Asymmetric Outer membrane | 70669 | 236 236 124 | 10 | 3 | 313 K |
| PDIM in MA | 64729 | 253 253 131 | 10 | 3 | 253 K, 313 K, 353 K |
| POPC only | 4725 | 87 87 85 | 3 | 3 | 313 K |
| PDIM in POPC | 50154 | 235 235 100 | 10 | 3 | 253 K, 313 K, 353 K |
| PDIM in PEPC | 46665 | 226 226 100 | 10 | 3 | 253 K, 313 K, 353 K |
| PDIM in PSPC | 64729 | 236 236 103 | 10 | 3 | 253 K, 313 K, 353 K |

[a]Number of replicas is 3 for any AA systems.

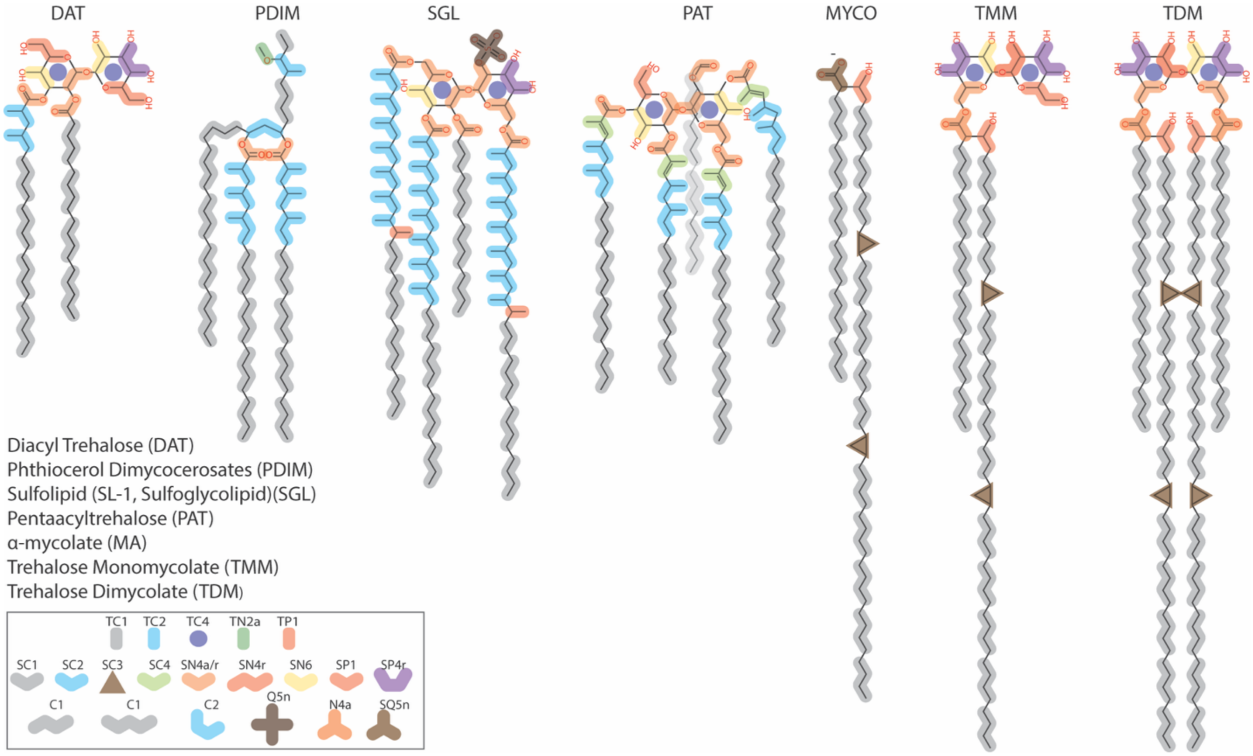


**Figure S1.** Atomic structures and corresponding coarse-grained (CG) mappings according to the MARTINI 3 force field. The lipids shown include diacyl trehalose (DAT), phthiocerol dimycocerosates (PDIM), sulfolipid-1 (SL-1; sulfoglycolipid, SGL), pentaacyl trehalose (PAT), α-mycolate (α-MA or MYCO), trehalose monomycolate (TMM), and trehalose dimycolate (TDM). The bead types used in the mapping are indicated in the bottom-right panel. TC4 beads are used to represent ring structures in lipid headgroups, and in trehalose-containing lipids, they correspond to the sugar headgroup. In contrast, PDIM lacks a sugar moiety, and an SC2 bead (blue) positioned above polar beads (orange) is used as the effective headgroup for analysis purposes. Charged beads are used as headgroups in SGL and α-MA, represented by Q5n and SQ5n bead types, respectively.

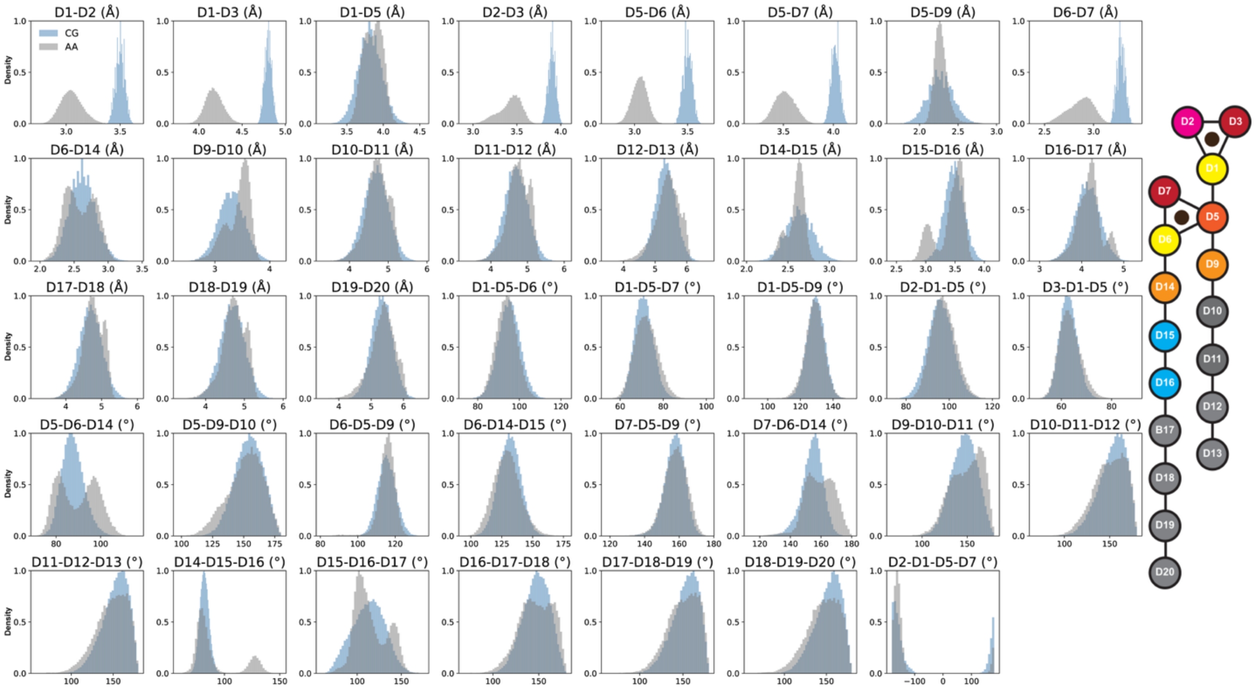


**Figure S2.** Comparison of the bonded parameters between AA and CG representations of DAT. The bonded distributions were calculated using all DAT molecules from all replicas of the asymmetric MOM (1X, 313 K) simulations, sampled at 1 ns interval over the time from 1 μs to 3 μs**.** The color of the beads corresponds the bead color scheme followed in the AA-CG mapping.

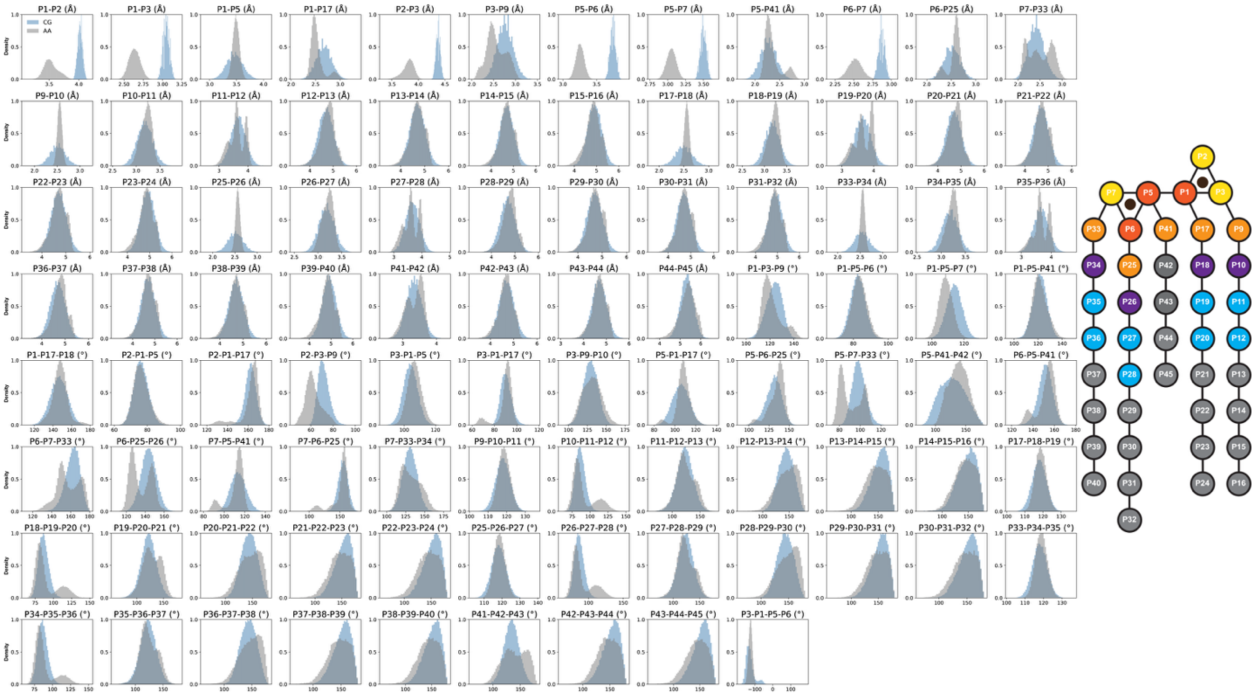


**Figure S3.** Comparison of the bonded parameters between AA and CG representations of PAT. The bonded distributions were calculated using all PAT molecules from all replicas of the asymmetric MOM (1X, 313 K) simulations, sampled at 1 ns interval over the time from 1 μs to 3 μs**.** The color of the beads corresponds the bead color scheme followed in the AA-CG mapping.

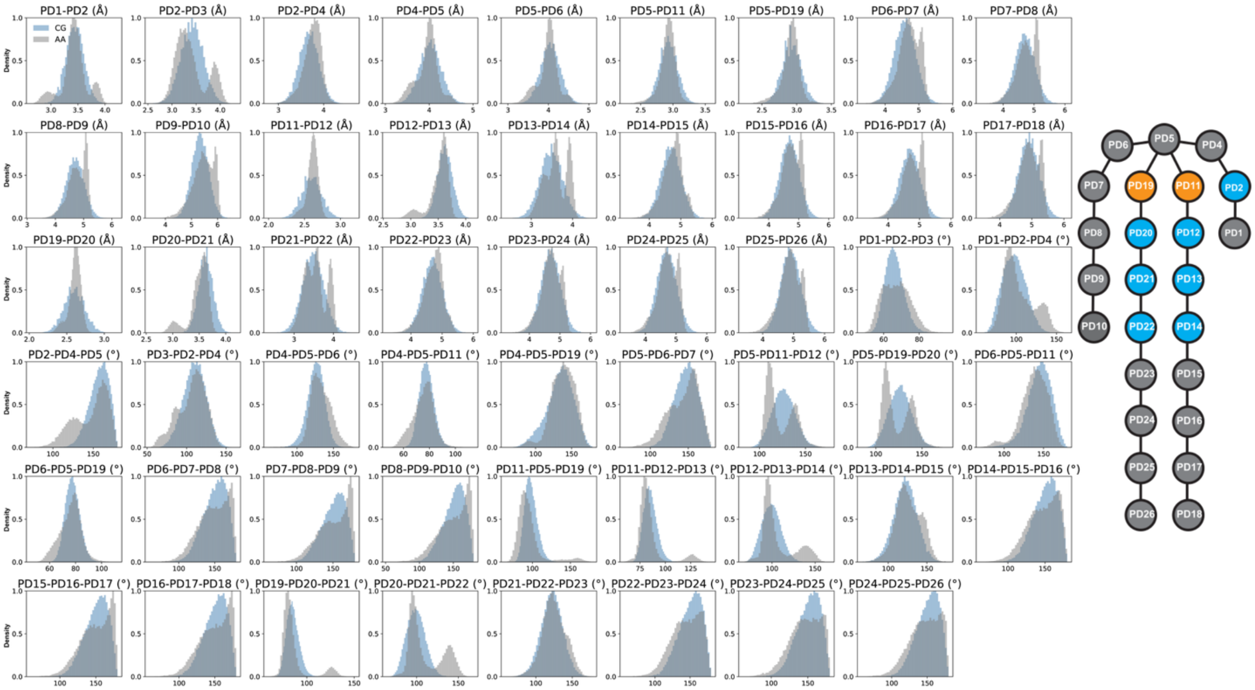

**Figure S4.** Comparison of the bonded parameters between AA and CG representations of PDIM. The bonded distributions were calculated using all PDIM molecules from all replicas of the asymmetric MOM (1X, 313 K) simulations, sampled at 1 ns interval over the time from 1 μs to 3 μs**.** The color of the beads corresponds the bead color scheme followed in the AA-CG mapping.

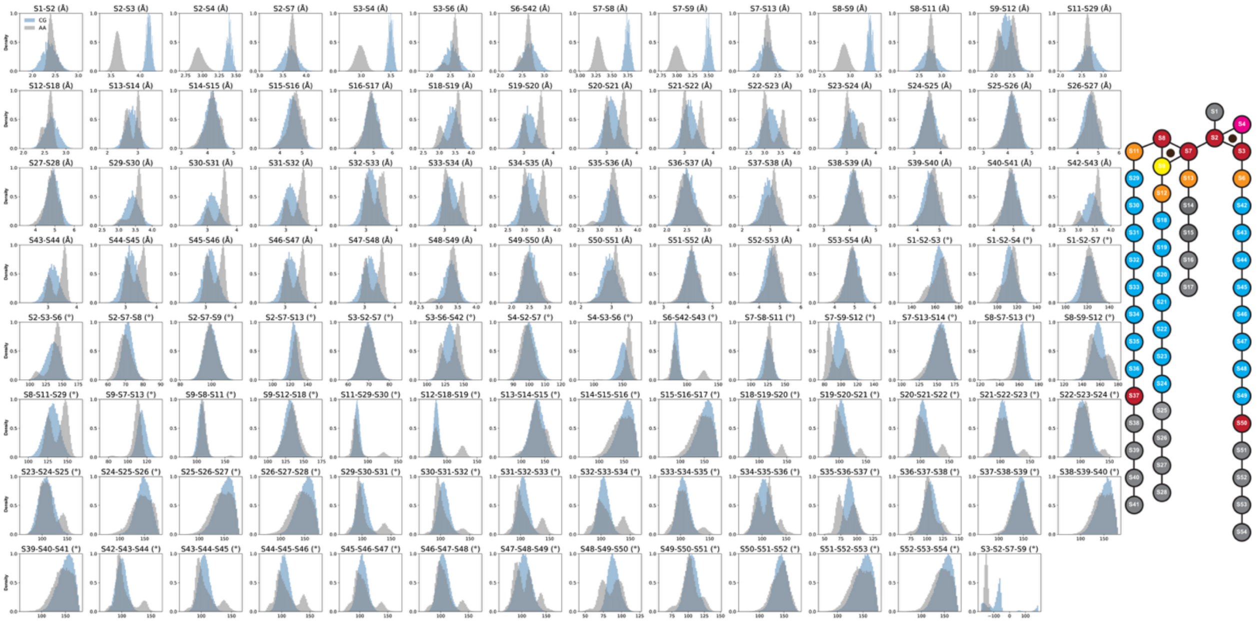

**Figure S5.** Comparison of the bonded parameters between AA and CG representations of SGL. The bonded distributions were calculated using all SGL molecules from all replicas of the asymmetric MOM (1X, 313 K) simulations, sampled at 1 ns interval over the time from 1 μs to 3 μs**.** The color of the beads corresponds the bead color scheme followed in the AA-CG mapping.

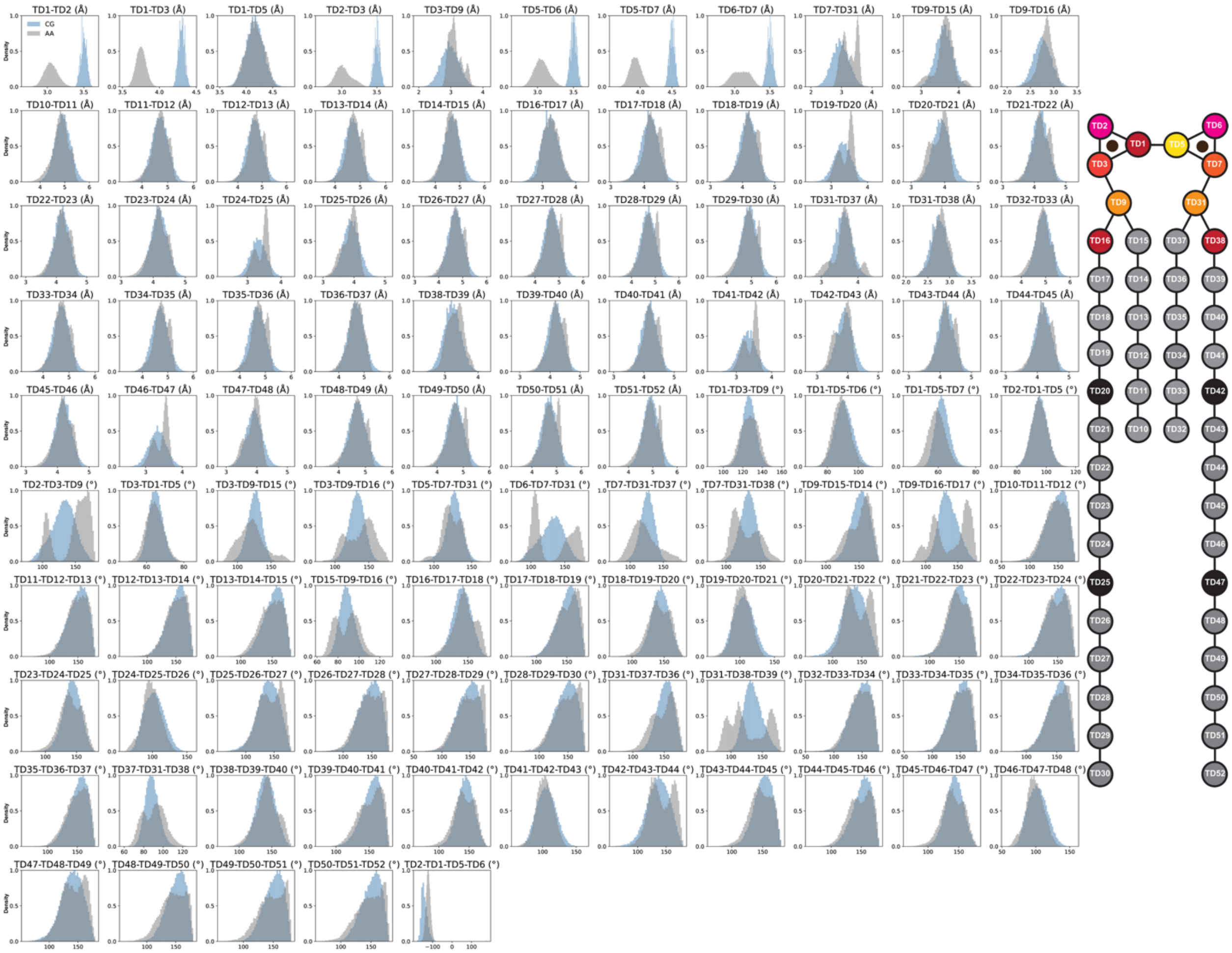

**Figure S6.** Comparison of the bonded parameters between AA and CG representations of TDM. The bonded distributions were calculated using all TDM molecules from all replicas of the asymmetric MOM (1X, 313 K) simulations, sampled at 1 ns interval over the time from 1 μs to 3 μs**.** The color of the beads corresponds the bead color scheme followed in the AA-CG mapping.

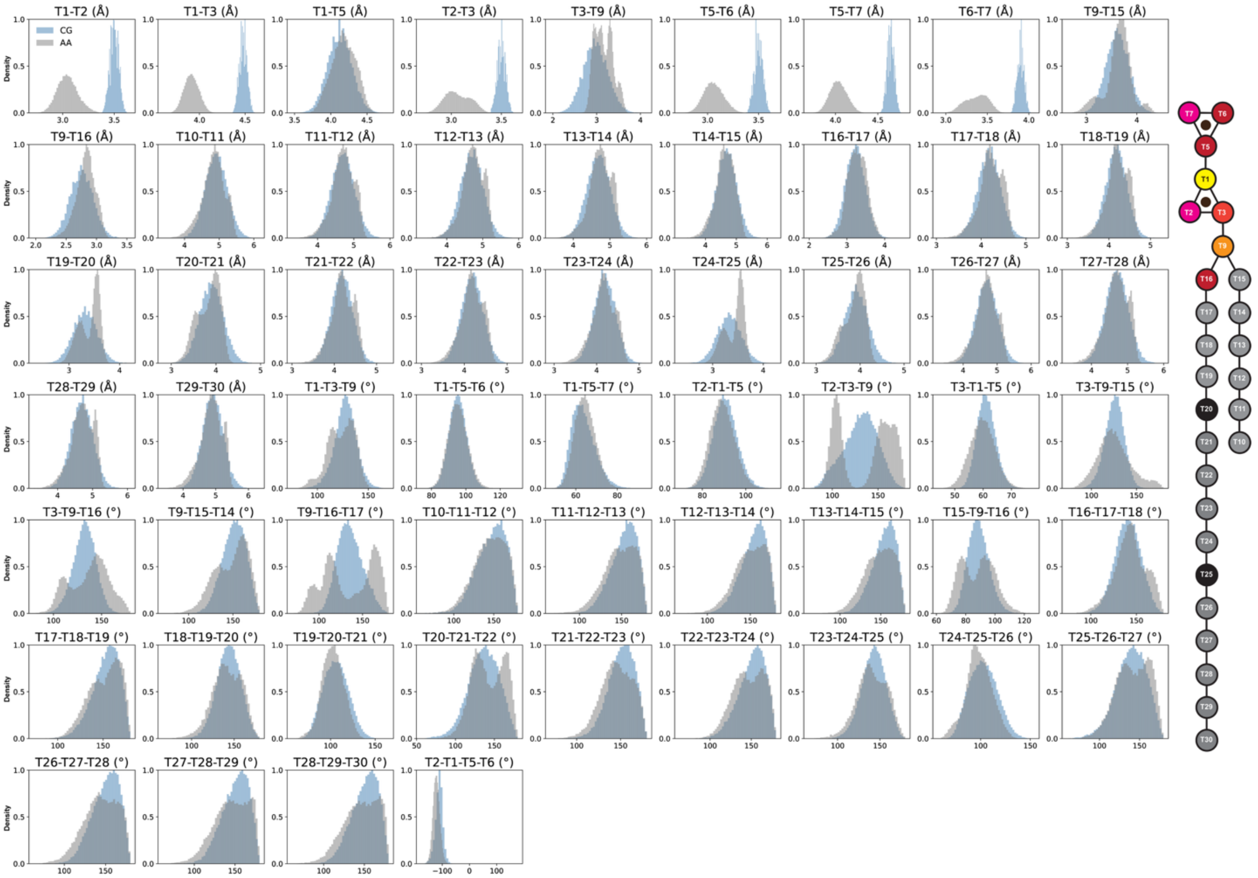


**Figure S7.** Comparison of the bonded parameters between AA and CG representations of TMM. The bonded distributions were calculated using all TMM molecules from all replicas of the asymmetric MOM (1X, 313 K) simulations, sampled at 1 ns interval over the time from 1 μs to 3 μs**.** The color of the beads corresponds the bead color scheme followed in the AA-CG mapping.

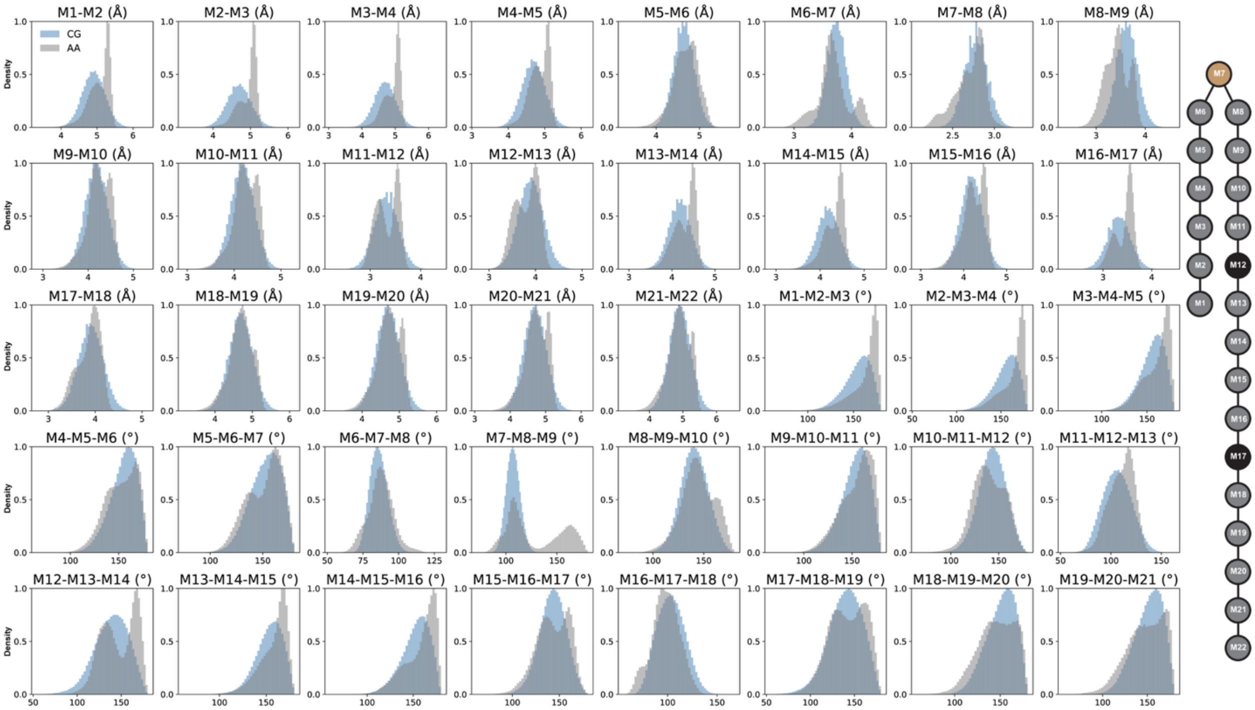


**Figure S8.** Comparison of the bonded parameters between AA and CG representations of α-MA. The bonded distributions were calculated using all α-MA molecules from all replicas of the asymmetric MOM (1X, 313 K) simulations, sampled at 1 ns interval over the time from 1 μs to 3 μs**.** The color of the beads corresponds the bead color scheme followed in the AA-CG mapping.

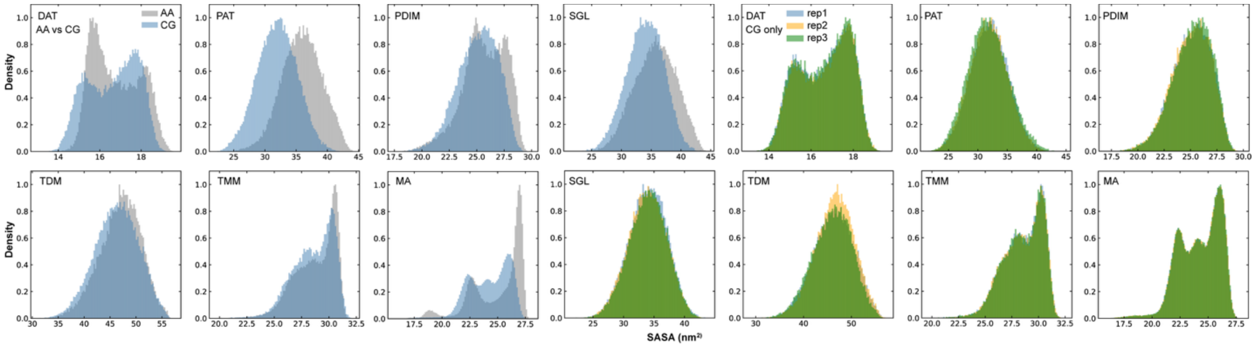


**Figure S9.** Solvent accessible surface area (SASA) comparison between AA (gray) and CG systems (light blue) at 313 K: data from 1 μs - 3 μs. The SASA profiles among various CG replicas are also shown in blue, orange, and green.

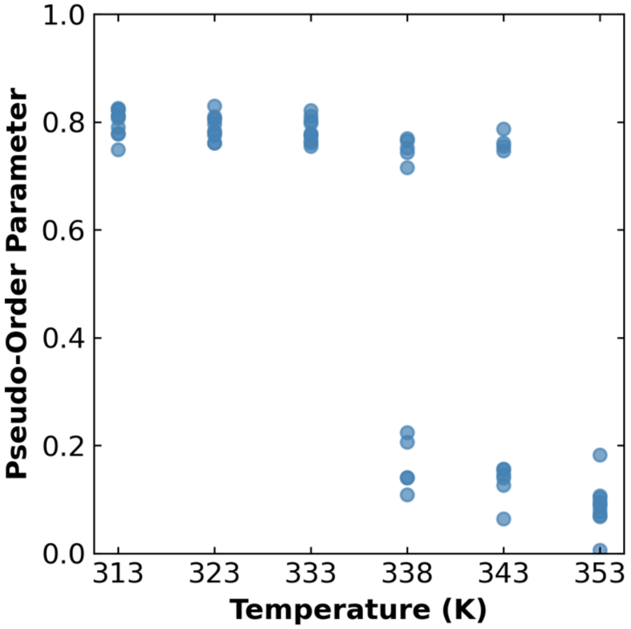


**Figure S10.** Pseudo-order parameters for individual CG replicas of the symmetric inner leaflet α-MA system. Each point represents a single replica at the indicated temperature. At 338 K and 343 K, variability in replica behavior is observed; some replicas retain high order, and others show a transition to a more disordered state, indicating heterogeneity in a phase transition among the replicas.

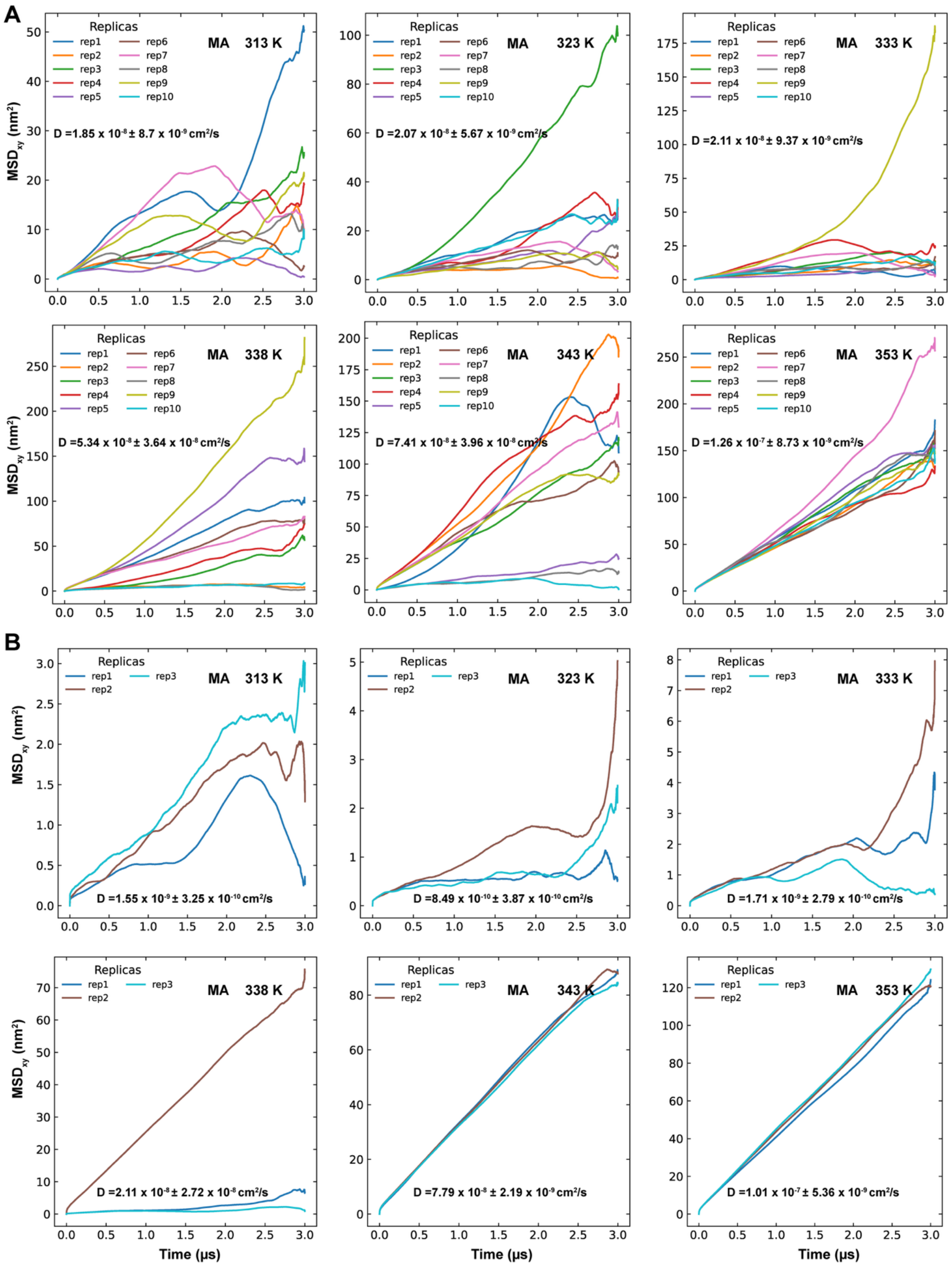
A
B
Replicas
rep1
rep2
rep3
rep4
rep5
rep6
rep7
rep8
rep9
rep10
MA 313 K
MA 323 K
MA 333 K
MA 338 K
MA 343 K
MA 353 K
D =1.85 x $10^{-8}$ ± 8.7 x $10^{-9}$ $cm^2/s$
D =2.07 x $10^{-8}$ ± 5.67 x $10^{-9}$ $cm^2/s$
D =2.11 x $10^{-8}$ ± 9.37 x $10^{-9}$ $cm^2/s$
D =5.34 x $10^{-8}$ ± 3.64 x $10^{-8}$ $cm^2/s$
D =7.41 x $10^{-8}$ ± 3.96 x $10^{-8}$ $cm^2/s$
D =1.26 x $10^{-7}$ ± 8.73 x $10^{-9}$ $cm^2/s$
D =1.55 x $10^{-9}$ ± 3.25 x $10^{-10}$ $cm^2/s$
D =8.49 x $10^{-10}$ ± 3.87 x $10^{-10}$ $cm^2/s$
D =1.71 x $10^{-9}$ ± 2.79 x $10^{-10}$ $cm^2/s$
D =2.11 x $10^{-8}$ ± 2.72 x $10^{-8}$ $cm^2/s$
D =7.79 x $10^{-8}$ ± 2.19 x $10^{-9}$ $cm^2/s$
D =1.01 x $10^{-7}$ ± 5.36 x $10^{-9}$ $cm^2/s$
$MSD_{xy}$ ($nm^2$)
Time (μs)

**Figure S11. A.** Mean squared displacement (MSD) profiles of α-MA of ten replicas in the symmetric inner leaflet systems at different temperatures (313 K, 323 K, 333 K, 338 K, 343 K, and 353 K). **B** MSD profiles of α-MA in three replicas of larger symmetric inner leaflet system at various temperatures. Diffusion coefficients are reported along with each graph; for the smaller systems diffusion coefficient were calculated using first 500 ns while for the larger system the diffusion coefficients were calculated using 200 ns - 1000 ns timeframe. The graphs show the noisy diffusion profiles for ordered phases (313 K, 323 K and 333 K), while linear profiles for disordered phase in both system sizes. The diffusion coefficients of α-MA in the larger system are notably smaller than that of smaller systems, but the diffusion coefficients of both system sizes remains comparable in the disordered phase.

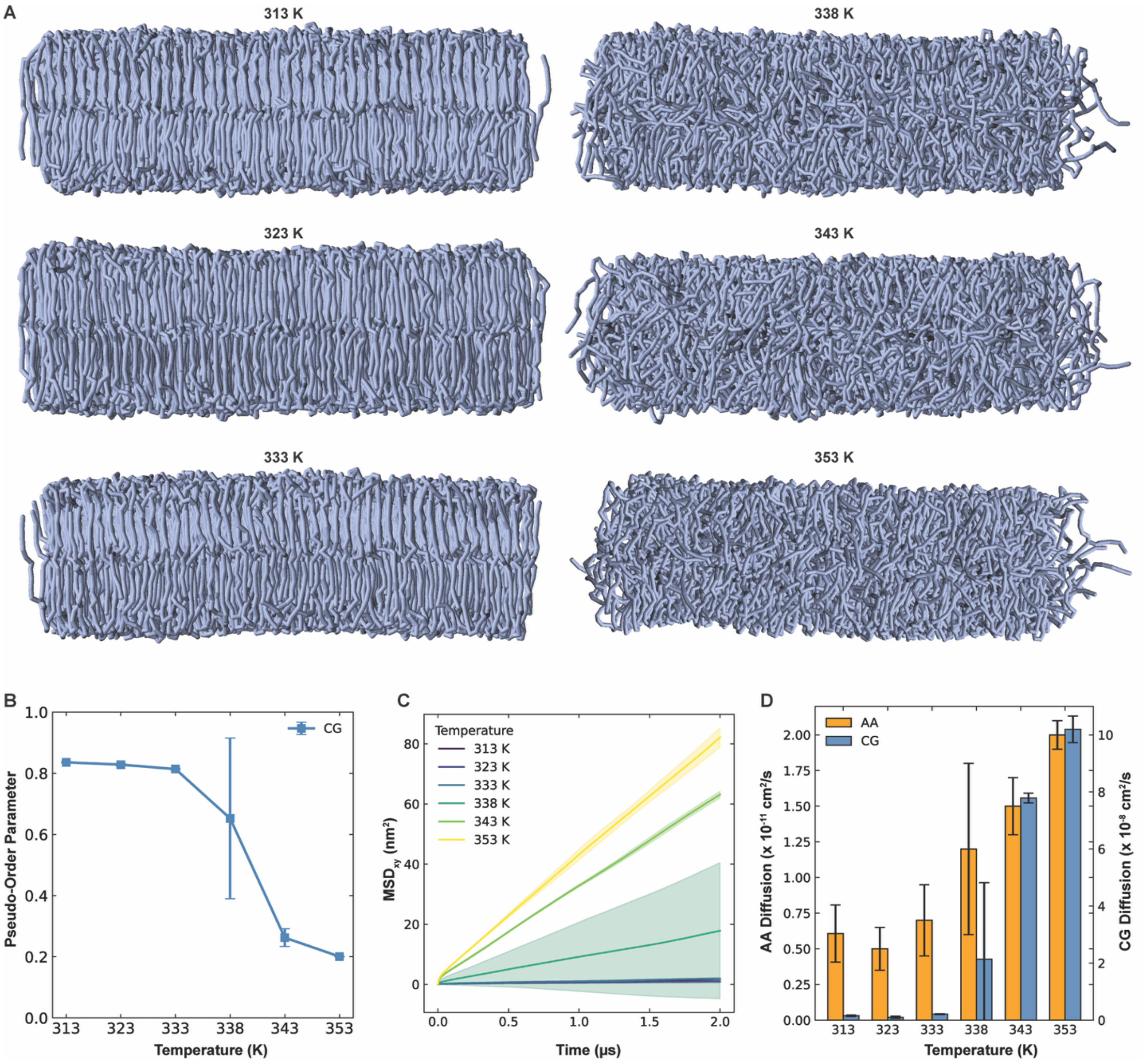


**Figure S12. A.** Snapshots (10 μs) of larger (9X) inner leaflet α-MA symmetric system at various temperatures (313 K, 323 K, 333 K, 338 K, 343 K and 353 K); a replica with disordered phase is shown for 338 K. **B.** Pseudo-order parameter for the lipid tails of α-MA in the larger system showing the gradual phase transition of the membrane at 338 K. The larger error bar at 338 K corresponds to ordered and disordered phase in three replicas. **C**. Mean-squared displacements (MSD) of α-MA in the larger system at various temperature, solid line represents the average diffusion, and shaded region represents the variation in the replicas. **D.** Diffusion coefficients of α-MA in the larger system compared to the diffusion coefficient of α-MA in the atomistic system showing the overestimation of the diffusion coefficient in CG systems. Diffusion coefficient is fitted between 200 ns - 1000 ns trajectory.

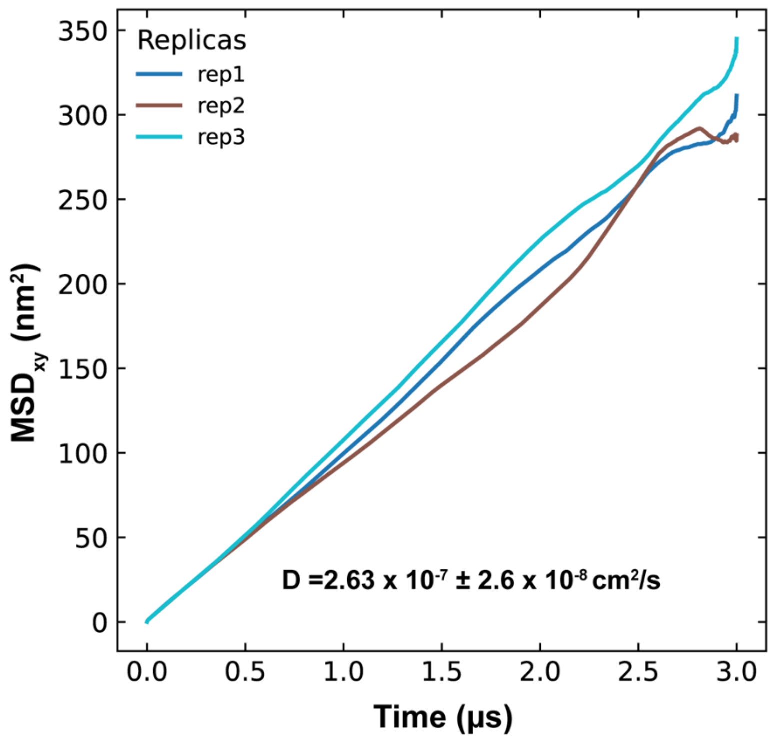


**Figure S13.** Diffusion coefficient of POPC calculated from POPC only system at 313 K. The diffusion coefficients were calculated using 200 ns - 1000 ns timeframe.

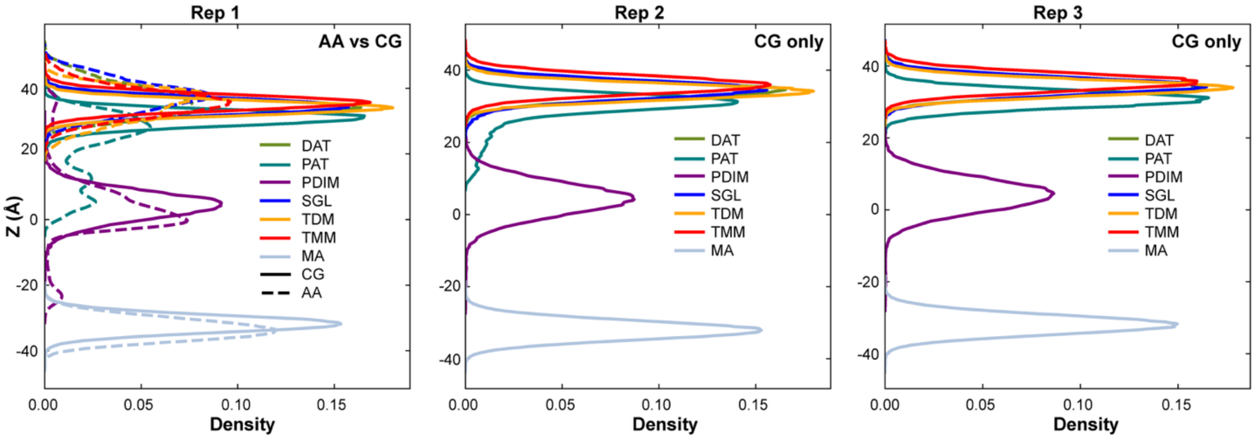


**Figure S14.** Lipid headgroup z-position densities (1 μs - 3 μs**)** in the asymmetric MOM CG (1X, 313 K) system across three replicas. In the first panel, a comparison with the AA system is also shown, while the remaining panels display only CG replicas individually.

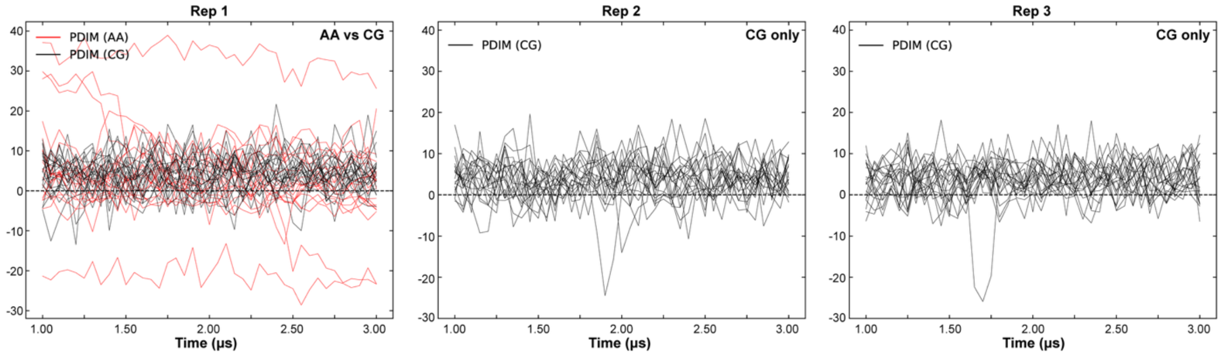


**Figure S15.** Timeseries (1 μs - 3 μs) of the z-position of the PDIM headgroup, showing its movement toward the center during the asymmetric MOM CG (1X, 313 K) simulations across three replicas. The first panel includes a comparison with the AA system, while the remaining panels show the CG replicas only.

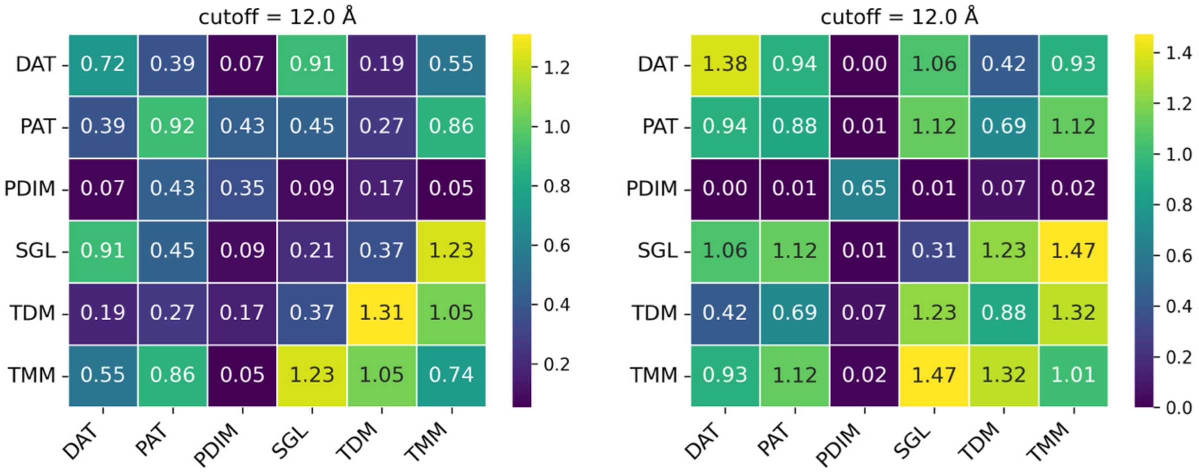


**Figure S16.** Average number of neighboring lipids within a 12 Å distance, calculated from the center of geometry of each lipid, showing no aggregation of individual lipids in both asymmetric MOM (1X, 313 K) AA (left) and CG (right) simulations. Data are averaged over three replicas from the trajectory between 1 μs and 3 μs**.**

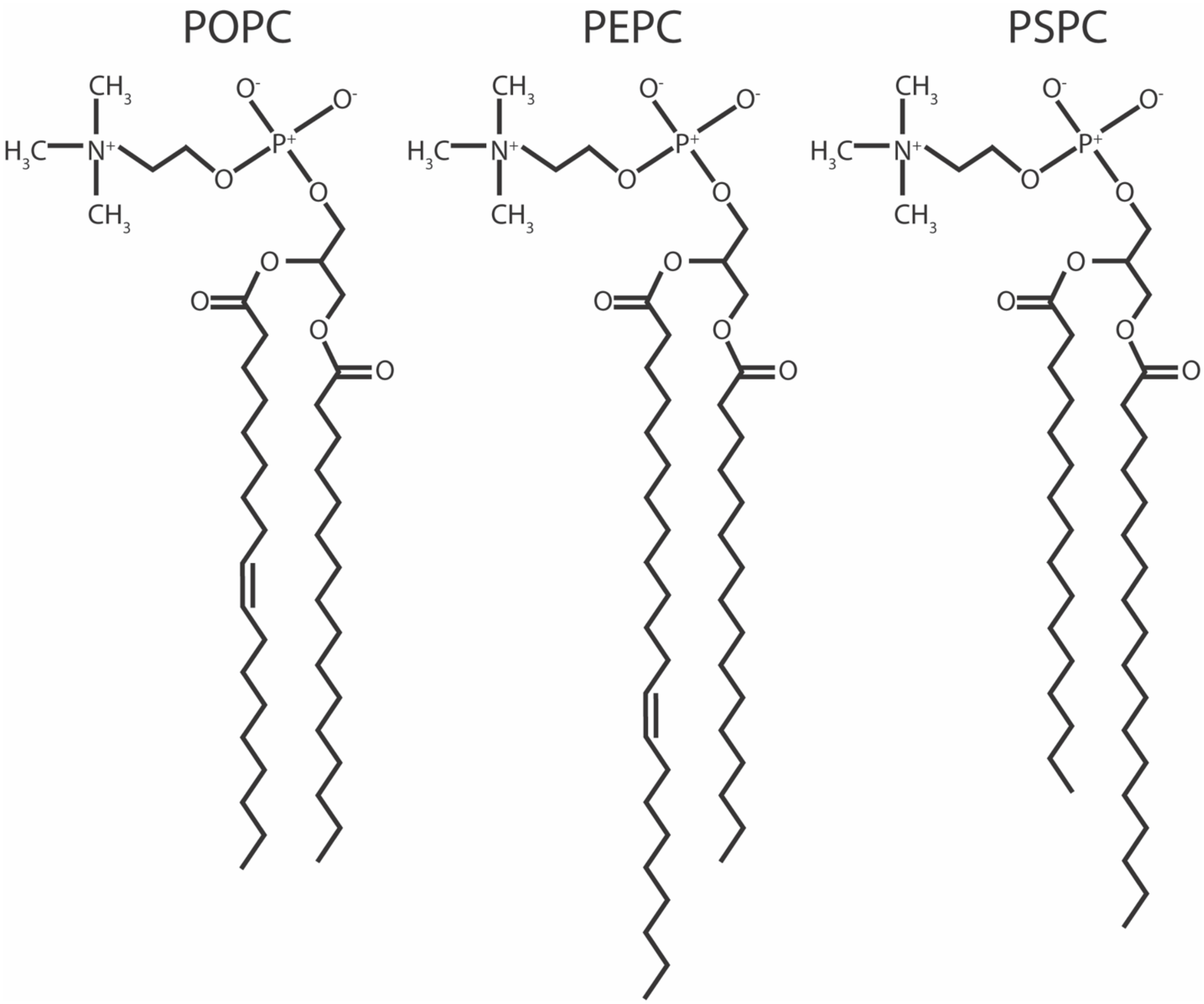


**Figure S17.** Chemical structures of POPC (palmitoyl oleoyl phosphocholine), PEPC (palmitoyl erucoyl phosphatidylcholine), and PSPC (palmitoyl stearoyl phosphocholine).

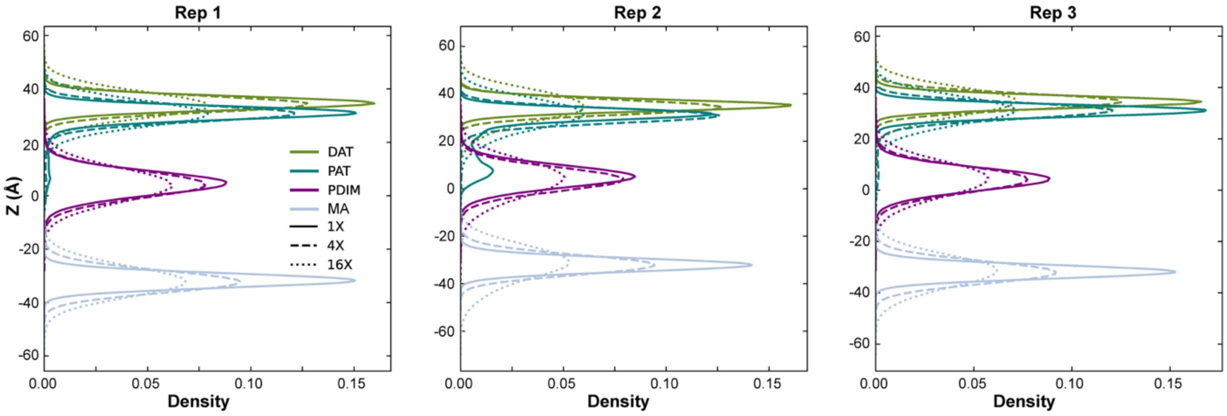


**Figure S18.** Lipid headgroup z-position densities across three replicas in systems of different lateral sizes (313 K): 1X (~100 Å × 100 Å, solid lines), 4X (~200 Å × 200 Å, dashed lines), and 16X (~400 Å × 400 Å, dotted lines). Data are collected at the interval of 1 ns over 1 μs – 3 μs.

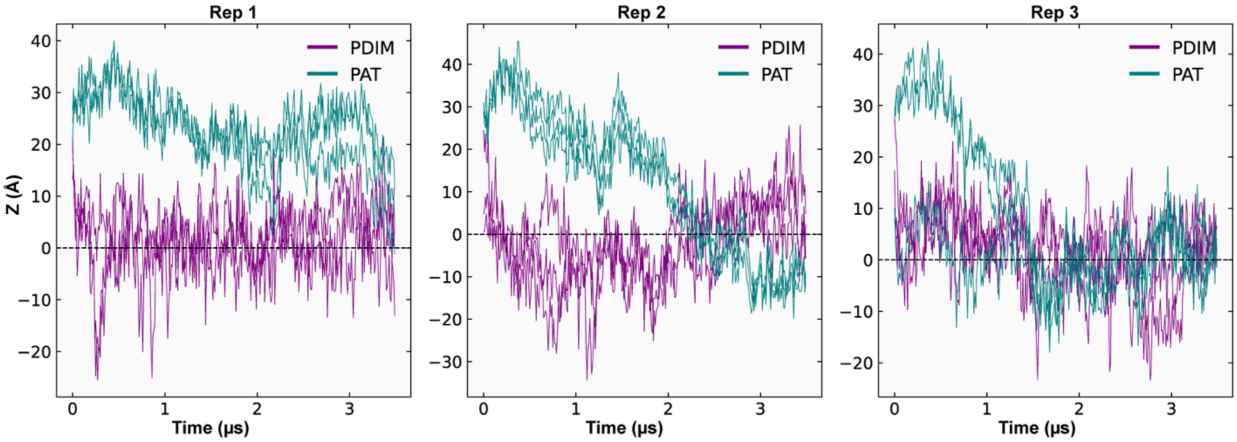


**Figure S19.** Timeseries of PAT and PDIM headgroups (showing the three lowest) illustrating the migration of PAT toward the center of the membrane in the 16X asymmetric MOM system (313 K), which was observed in smaller AA system simulations, but not in smaller CG system simulations.

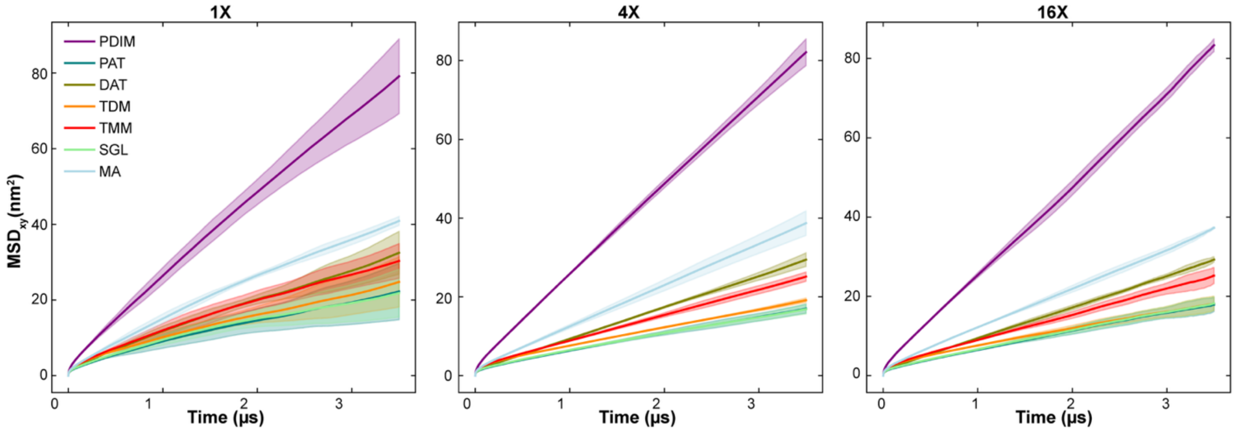


**Figure S20.** Lateral MSD plots (0 μs – 3.5 μs) for various lipids in the asymmetric MOM (313 K) simulations over three different system sizescales. The mobility remains more or less consistent regardless of size, with PDIM and α-MA being the most diffusive, while larger and branched lipids show slower diffusion.

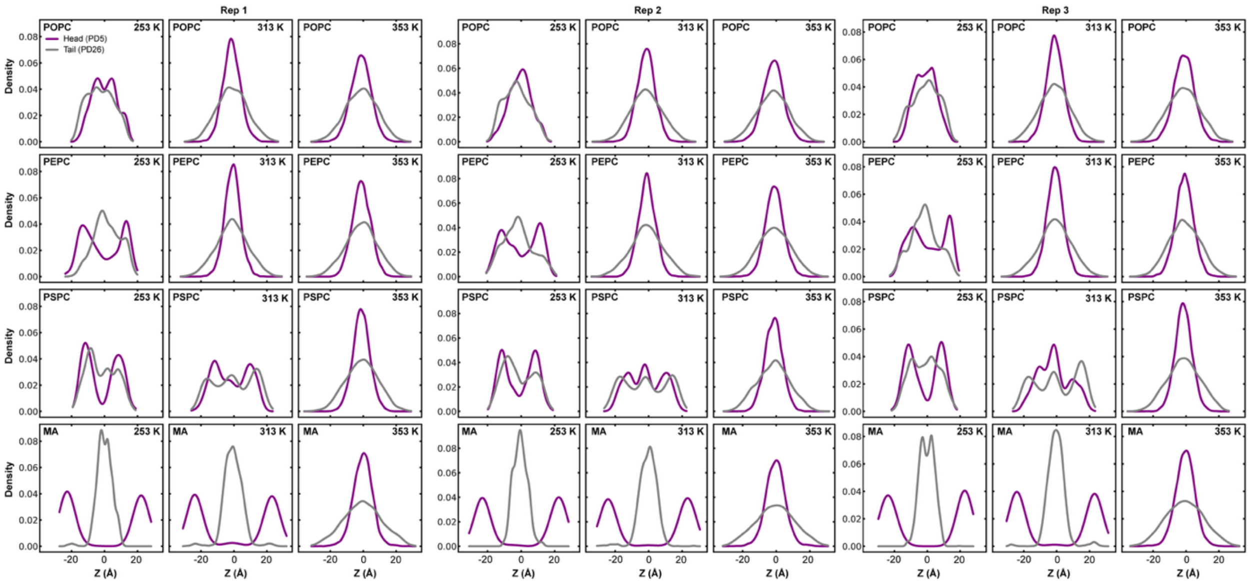


**Figure S21.** Z-density profiles of PDIM headgroup (PD5, purple) and tail (PD26, grey) beads across three independent replicas at varying temperatures (253 K, 313 K, and 353 K). In all replicas, increasing temperature leads to a consistent shift of the PDIM headgroup from the membrane surface toward the membrane center, reflecting enhanced membrane fluidity.

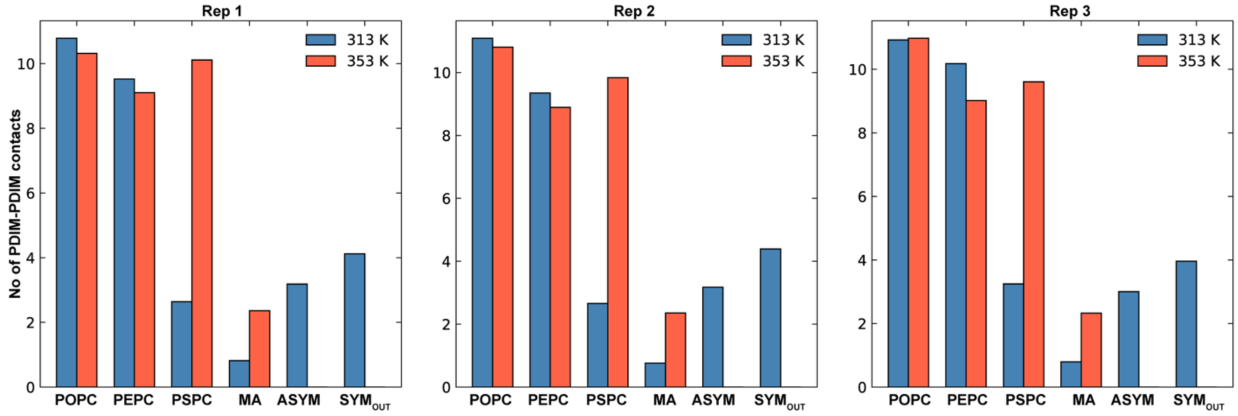


**Figure S22.** Comparison of molecular contacts (within 6 Å) at 313 K and 353 K. The results across three replicates consistently show increased PDIM aggregation in fluid membrane phases.

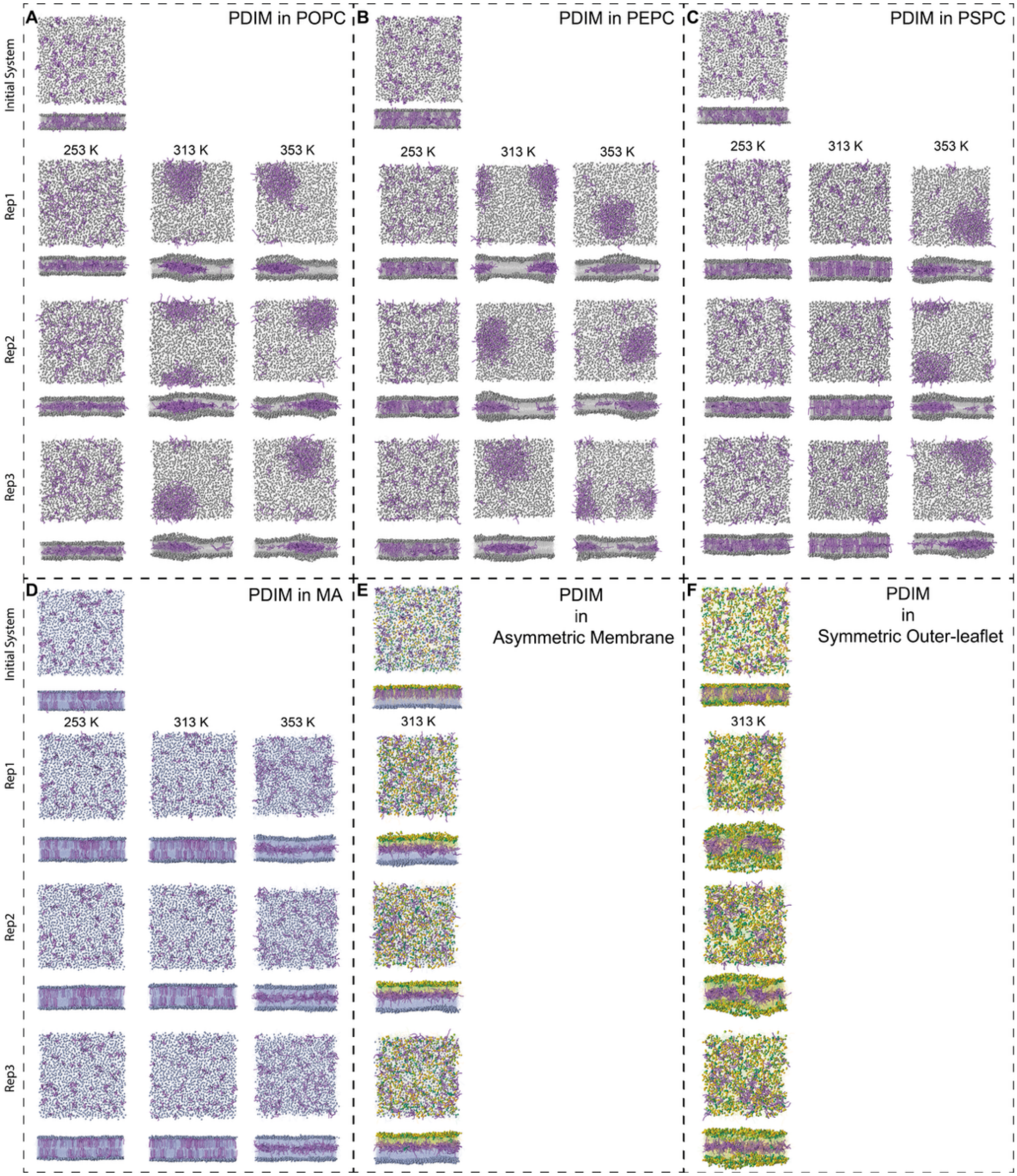


**Figure S23.** PDIM behavior in different lipid environments. All snapshots were taken at 5 μs, except for the initial system and the symmetric outer leaflet system, for which 2 μs snapshots are shown due to membrane instability. Both the lateral and top views are shown. In POPC, PDIM migrates toward the membrane center at all temperatures studied, although aggregation is not observed at 253 K. In PEPC, fewer PDIM molecules migrate toward the membrane center at 253 K, whereas at higher temperatures PDIM migrates to the center and forms aggregates. In PSPC, PDIM does not migrate toward the membrane center at 313 K, but both migration and aggregation are observed at 353 K. In α-MA, PDIM migrates toward the membrane center at 353 K without forming aggregates. In the asymmetric outer leaflet system, PDIM migrates toward the membrane center at 313 K without aggregation. In contrast, in the symmetric outer leaflet system, PDIM also migrates toward the membrane center at 313 K, but membrane disruption occurred at longer timescale, and no aggregation is observed. Overall, these observations demonstrate that PDIM translocation across the membrane and its aggregation are strongly dependent on membrane fluidity.
